# Breaking Barriers: pCF10 Type 4 Secretion System relies on a self-regulating muramidase to modulate the cell wall

**DOI:** 10.1101/2024.02.15.580431

**Authors:** Wei-Sheng Sun, Gabriel Torrens, Josy ter Beek, Felipe Cava, Ronnie P-A Berntsson

## Abstract

Conjugative Type 4 Secretion Systems (T4SS) are a main driver for the spread of antibiotic resistance genes and virulence factors in bacteria. To deliver the DNA substrate to recipient cells, it must cross the cell envelopes of both donor and recipient mating bacteria. In the T4SS from the enterococcal conjugative plasmid pCF10, PrgK is known to be the active cell wall degrading enzyme. It has 3 predicted extracellular hydrolase domains, LytM, SLT and CHAP. Here, we report the structure of the LytM domain, and show that its active site is degenerate and lacks the active site metal. Further, we show that only the predicted SLT domain is functional, and that it unexpectedly has a muramidase instead of a lytic transglycosylase activity. While we did not observe any peptidoglycan hydrolytic activity for the LytM or CHAP domain, we found that these domains downregulated the SLT muramidase activity. The CHAP domain was also found to be involved in PrgK dimer formation. Further, we show that PrgK interacts with PrgL, which likely targets PrgK to the rest of the T4SS. The presented data provides important information towards understanding the function of Gram-positive T4SSs.

## Introduction

Enterococci are gram-positive (G^+^) bacteria commonly found in the environment and in animal microbiomes. Enterococci are also opportunistic pathogens that are prevalent in hospital acquired infections^1–3^. These bacteria efficiently form biofilms^4,5^, and are often resistant to multiple antibiotics, including last resort antibiotics such as vancomycin and linezolid^6,7^. They are also highly adept in transferring their resistance to other bacteria via horizontal gene transfer, usually via conjugation. Conjugation is mediated by Type 4 Secretion Systems (T4SSs) that transfer DNA and protein(s) from a bacterial donor to a recipient cell. T4SSs are multi-protein complexes that span across the bacterial cell membrane(s) and cell wall. During the past years, our knowledge of T4SSs has greatly expanded, with published structures of the Dot/Icm system from *Legionella pneumophila*, the Cag system from *Helicobacter pylori,* and the T4SSs from the pKM101, R388, and F plasmids (all from *E. coli*)^8–12^. For a recent review of G^-^ T4SSs and their structure and function, please see Costa *et al, 2023*^13^. The best-known G^+^ T4SSs are encoded on the following plasmids: pCF10 and pIP501 from *E. faecalis*, and pCW3 from *Clostridium perfringens*^14–16^. Among these, the T4SS from the single, tightly regulated, operon on pCF10 is likely the best characterized^14,17–19^. We and others have previously characterized the adhesin proteins that facilitate mating pair formation^20–22^, as well as the DNA transfer and replication proteins^23–25^. However, compared to the information from G^-^ T4SS systems, we still have only a very limited insight into G^+^ T4SSs^15^.

One of the major differences between G^-^ and G^+^ bacteria is the thickness of the peptidoglycan (PG) layer that makes up the cell wall. While G^-^ cell walls are usually only a few nanometers thick (up to 4 nm in *E. coli*)^26^, the cell-walls of G^+^ bacteria are 30-100 nanometers^27^. The cell-wall of *E. faecalis* has been reported to be ca. 40 nm thick^28^. In all cases, this cell-wall forms a barrier that must be crossed during conjugation. In the characterized G^-^ systems the T4SSs span over the inner membrane, the periplasmic space (including the PG) and the outer membrane. In G^+^ systems this is predicted to be very different, with the T4SS channel spanning the single membrane but not through the entire PG layer. It is, therefore, not surprising that the PG-degrading enzymes that create specific lesions in the cell wall differ between G^-^ and G^+^ T4SSs. In G^-^ systems, the PG-degrading enzymes are usually soluble proteins that are secreted to the periplasm and needed to create lesions in the PG to allow for the T4SS channel assembly^29–31^. In G^+^ systems, these enzymes are usually anchored to the cell membrane via a transmembrane helix and are thought to be required to create defined lesions in the PG layer to allow substrate transfer^32,33^. In T4SSs, they are divided into 4 classes, ALPHA, BETA and DELTA, which are found in G^+^ T4SS, and the GAMMA class, which is present in G^-^ T4SSs^34^. The pCF10 encoded PrgK is essential for conjugation and belongs to the DELTA family of larger PG-degrading enzymes^34^. Previous work has indicated that PrgK consists of an intracellular N-terminal domain, a transmembrane helix and an extracellular part consisting of a LytM (metallo-peptidase), an SLT (a Soluble Lytic Transglycosylase) and a CHAP (Cysteine, Histidine-dependent Amidohydrolases/Peptidases) domain^32^. While all 3 extracellular domains can bind PG *in vitro* and their over-expression led to cell-morphology defects in *E. coli*, PG hydrolysis activity could only be measured for the SLT domain^32^. However, PrgK variants lacking the SLT domain were reported to partially complement a *prgK* knock-out strain *in vivo*, indicating that at least one other domain retains hydrolysis activity. It has been unclear how PrgK is regulated or how it interacts with the remaining T4SS channel proteins.

Here, we report the crystal structure of the LytM domain and combine AlphaFold models with biochemistry to analyze the remaining part of the extracellular domains. Alphafold modelling suggested that the extracellular domain of PrgK interacts with that of PrgL, another pCF10 encoded T4SS protein that plays a role in T4SS biogenesis^35^. We confirm this interaction *in vitro* and show that PrgK can also dimerize in a redox dependent manner. Finally, we characterized PrgK activity *in vitro* and revealed that the predicted SLT domain exhibits muramidase instead of the previously predicted lytic transglycosylase activity, and that this domain is negatively regulated by the other two extracellular domains: LytM and CHAP.

## Materials and methods

### Strains and Culture Conditions

See Table S1 for detailed plasmid and strain information. In brief, *E. coli* Top10 was the host strain for plasmid construction and propagation, and *E. coli* BL21(DE3) was the host strain for IPTG-inducible protein overexpression. *E. coli* strains were cultivated in Luria Bertani (LB) broth with necessary selection antibiotics at 37 °C with agitation. *E. faecalis* strains were cultured with Brain-Heart Infusion (BHI) for overnight incubation or in Tryptic soy broth without dextrose (TSB-D) with antibiotics required for plasmid selection. Concentrations of antibiotics for *E. coli* selection were as follows: kanamycin (50 μg/ml), chloramphenicol (25 μg/ml). In *E. faecalis* cultures, antibiotics were used as the following concentrations: tetracycline (10 μg/ml), fusidic acid (25 μg/ml), erythromycin (100 μg/ml), spectinomycin (1000 μg/ml). Plasmids transformations to *E. coli* were carried out by standard heat-shock, whereas *E. faecalis* strains were transformed by electroporation^36^.

Overnight cultures of *E. faecalis* OG1RF transformed with empty pMSP3545S nisin-inducible vector or with the same vector harboring *prgK* ^37^, were inoculated in fresh TSB-D medium with selection antibiotic (erythromycin) and 50 ng/mL nisin to an initial OD600nm of 0.05. After incubating at 37°C with agitation for the indicated time (1, 1.5, 2, 2.5, 3.5 and 5 hrs), optical density was measured again for growth curve. For *E. faecalis* cell viability, cell cultures were harvested after 3.5 hours. Individual samples were serial diluted and then plated out on BHI agar plates. The resulting plates were incubated at 37°C for colony forming unit enumeration.

### Scanning Electron Microscopy

*E. faecalis* cells for scanning EM were cultured and induced as described in the previous paragraph. After 2 or 3.5 hrs incubation, 5 mL of the cultures was spun down at 3000 *×g*, 4 °C for 10 min. Pellets were resuspended in 400 µL 0.1 M Na-Phosphate buffer pH 7.4 and transferred to a 1.5 mL tube. The cells were again pelleted, but now at 20,000 *×g*, 4 °C, 10 min. All supernatant was removed, and the pellets were carefully resuspended in 200 µL 0.1 M Na-Phosphate buffer pH 7.4 (by both pipetting and vortexing). Afterwards 200 µL 5% Gluteraldehyde in 0.1 M Na-Phosphate buffer pH 7.4 was added, mixed by pipetting up and down to crosslink the cells and incubated for several hours at 4°C. The mixtures were then further incubated overnight at 4°C on top of polylysin coated coverslips that were placed at the bottom of a 24-well plate. Samples were washed 3 times with 0.1M phosphate buffer and then dehydrated with graded series (50%, 70%, 80%, 90%, 100%) of ethanol. They were further critical point dried with Leica EM300. The samples were coated with a 5 nm layer of Pt (Quorom Q150T ES) and imaged using a Field emission Scanning Electron Microscope (FESEM, Carl Zeisss Merlin) at an accelerating voltage of 5kV and probe current of 120 pA.

### Cloning and plasmids

To characterize the extracytoplasmic region of PrgK, DNA sequences covering different fragments of PrgK were amplified for protein expression, and subsequent studies. Domain boundaries were identified using Phyre2 and RaptorX^38,39^. pMSP3545-prgK^32^ was used as template along with the corresponding primer pairs (Table S1) to amplify PCR products covering sequences coding for PrgK_EC_ (residues 273-871), PrgK_SLT-CHAP_ (residues 530-871), PrgK_LytM_ (residues 273-529), PrgK_SLT_ (residues 539-723), and PrgK_CHAP_ (residues 723-871); and pMSP3545-prgK:_C766A H828A_ was the template for generating amplicons for PrgK_EC: C766A H828A_ and PrgK_SLT-CHAP: C766A H828A_. These PCR products were cloned into pINIT_Kan vector for sequencing and then sub-cloned to p7X expression vectors (p7XC3GH and p7XNH3) following the guidelines of the FX-cloning system^40^. p7XNH3-prgK_EC:C766A_ was constructed by the same site-directed mutagenesis protocol as previously described^20^ with partial-overlapping primer pair PrgK_C766A_inv-F and PrgK_C766A_inv-R.

### Protein expression and purification

p7X expression plasmids harboring the His-tagged PrgK variants-encoding sequences were transformed to *E. coli* BL21(DE3). The transformants were firstly inoculated in LB broth for small scale overnight incubation (37°C, 200 rpm), then diluted 1:100 in Terrific broth (TB) and further incubated at 37°C in the LEX bioreactor system (Epiphyte3). When the optical density reached 1.5, the incubation temperature was lowered to 18 °C and expression was induced by adding 0.1 mg/ml IPTG for 18 h. Afterwards, cells were pelleted, resuspended in lysis buffer (50 mM Na2HPO4/NaH2PO4, 300 mM NaCl, 10 mM imidazole, pH 7.0) in a ratio of 7g/100 ml and lysed at 4 °C and 25 kPsi using a Cell Disruptor (Constant System). Cell debris was removed by centrifugation for 30 minutes at 13,000 × *g*, 4°C and the supernatant was incubated at 4 °C with 1.25 ml Ni-NTA agarose resin (Macherey-Nagel) for 1 h with agitation. By gravity, the Ni-NTA resin was washed with 40 column volumes of wash buffer (50 mM Na2HPO4/NaH2PO4, 300 mM NaCl, 50 mM imidazole, pH 7.0). The bound proteins were eluted from the column by incubation with PreScission protease in a 1:100 ratio for 20 h at 4 °C, thereby removing the histidine affinity tag. Tag-free proteins were then loaded on a Superdex-200 Increase 10/300 GL column (Cytiva) equilibrated in one of the following buffers. Buffer for PrgK_LytM_ was 50 mM HEPES, 500 mM NaCl, pH 7.5; for PrgK_EC_ and PrgL32-208 mixtures. For the individual fragments of PrgK_EC_, PrgK_SLT-CHAP_, PrgK_SLT_, Prg_KCHAP_, a buffer of 100 mM citrate-citric acid, pH 5.0, 100 mM NaCl was used.

### Crystallization and structure determination

SEC-purified PrgK_LytM_ in 50 mM HEPES, 500 mM NaCl, pH 7.5, with a protein concentration of 15 mg/mL, was used for crystallization trials. Crystals were observed after 3-5 days, at 20 °C by sitting drop vapor diffusion in a condition with 0.2 M Ammonium sulfate, 0.1 M Sodium acetate, pH 4.6, 30 % w/v PEG 2000 MME and a protein to reservoir ratio of 1:1 in the drop. Crystals were flash cooled in liquid nitrogen. X-ray diffraction data was collected on beamline ID30B at the ESRF, France. The data was processed using XDS^41^. The Prg_KLytM_ crystals belongs to the space group of P1 and contain 2 molecules in the asymmetric unit. The crystallographic phase problem was solved by molecular replacement of PHASER ^42^, using the Alphafold2 model of LytM (residue 273-529) as a search model. Further model refinement was done by using PHENIX refine^43^ and further modelled using COOT^44^. The structure was refined to 1.5 Å with crystallographic Rwork/Rfree values of 17.7/20.8% (Table 1). The final Prg_KLytM_ model consists of residues 274-529, and was validated using MolProbity^45^. Atomic coordinates and structure factors have been deposited in the Protein Data Bank (PDB Code: 8S0U).

**Table 1.** Data collection and refinement statistics.

| **Data collection summary** | **LytM** |
| --- | --- |
| Resolution range | 49.13 – 1.5 (1.554-1.5) |
| Space group | P1 |
| Cell dimensions |  |
| a, b, c (Å) | 53.32, 53.69, 56.35 |
| $\alpha$, $\beta$, $\gamma$ (°) | 63.3, 70.4, 70.2 |
| Total reflections | 144756 (14115) |
| Unique reflections | 76670 (7437) |
| Multiplicity | 1.9 (1.9) |
| Completeness (%) | 93.35 (90.66) |
| Mean I/sigma (I) | 6.98 (2.9) |
| R-meas | 0.12 (0.42) |
| CC(1/2)* | 0.977 (0.942) |
| **Refinement summary** |  |
| R-work | 0.177 (0.259) |
| R-free | 0.208 (0.289) |
| Number of non-hydrogen atoms | 4961 |
| protein | 4268 |
| solvent | 693 |
| RMS(bonds) | 0.012 |
| RMS(angles) | 1.09 |
| Ramachandran favored (%) | 97.45 |
| Ramachandran allowed (%) | 2.55 |
| Ramachandran outliers (%) | 0.00 |
| Average B-factor | 24.94 |
| macromolecules | 23.30 |
| solvent | 35.02 |

Statistics for the highest-resolution shell are shown in parentheses.

### AlphaFold modelling

ColabFold^46^ (v1.5.2) was used to generate predictions of full-length PrgK using default settings, whereas the prediction of PrgK-PrgL interaction was carried out with AlphaFold-Multimer using default settings.

### Size exclusion chromatography with multi-angle static light scattering (SEC-MALS)

For analysis of the PrgK_EC_ dimer, 150-300 µL 9.6 mg/mL PrgK_EC_ (with a theoretical mass of 67.4 kDa) was loaded on a Superdex 200 Increase 10/300 GL column, equilibrated in SEC buffer (100 mM citrate-citric acid, pH 5.0, 100 mM NaCl) via an ÄKTA Pure (Cytiva) that was coupled to a light scattering (Wyatt Treas II) and refractive index (Wyatt Optilab T-Rex) detector to determine the molecular weight of the two eluting components (at ∼12 and ∼14.2 mL) via multi-angle laser light scattering (SEC-MALS).

For analysis of the PrgK-PrgL complex, 3-4 mg/mL PrgK_EC_ was mixed with PrgL32-208 in a molar ratio of 1:1 or 1:3 and incubated for 15 min at 4 °C. For three analyses, 200 uL of the 1:1 mixture was loaded directly on a Superdex 200 Increase 10/300 GL column, equilibrated in SEC buffer (100 mM citrate-citric acid, pH 5.0, 100 mM NaCl) via an ÄKTA Pure (Cytiva) that was coupled to a light scattering (Wyatt Treas II) and refractive index (Wyatt Optilab T-Rex) detector to determine the molecular weight of the two eluting components (at ∼12.5 and ∼13.8 mL) via multi-angle laser light scattering (SEC-MALS). For a fourth analysis the same was done, but then with the 1: 3 mixture. For four additional analyses a preparative SEC run on a similar column (Superdex 200 Increase 10/300 GL) in the same SEC buffer with an Äkta Purifier (Cytiva) was done prior to the SEC-MALS analysis to isolate the earlier eluting peak fraction (∼12.5 mL). Data was analyzed using Astra software (version 7.2.2; Wyatt Technology).

### Sacculi preparation

An established procedure was adopted^47^, albeit with significant changes, to purify the PG of the strains under investigation. In summary, 200 mL cultures were grown to an OD595nm of 0.5, and cells were harvested at 3,000 *×g* for 15 min. Pellets were boiled with magnetic stirring with 5% SDS for 3 hours and then left overnight stirring at RT. Subsequently, the sacculi was pelleted by ultracentifugation for 10 min at 20 °C and 150,000 *×g* and resuspended in 3 mL H2O. This was repeated three times to remove all SDS.

### E. faecalis peptidoglycan isolation

To isolate PG from *E. faecalis*, SDS-free pellets from sacculi preparation were resuspended in 1 mL Tris-HCl 100 mM pH 7.5 and then glass beads measuring 0.1 mm in diameter were added (200 mg) and subjected to 10 min of vortexing at 4°C. The samples were subjected to a short spin (1 min) at 2,000 *×g* to allow precipitation of the glass beads and debris. Carefully, the supernatant was recovered and ultracentrifuged at 150,000 *× g* for 10 min at 20 °C. After centrifugation, the pellets were resuspended in 1 mL Tris-HCl, 100 mM pH 7.5 and subjected during 2 hours of treatment with 40 µL 1 M MgSO4, 2 µL RNase A (500 µg mL^−1^), and 1 µL DNase I (100 µg mL^−1^), followed by an incubation with 50 µL CaCl2 and 100 µL trypsin (2 mg mL^−1^) for 16 hours with magnetic stirring. Digestion was inactivated by adding 200 µL SDS 10% (w/v) and boiled for 10 min. Samples were centrifuged at 150,000 *× g* for 10 minutes at 20 °C and washed with MilliQ water each time until the SDS was completely removed. SDS-free pellets were treated with 1 mL of 8M LiCl for 10 min at 37 °C and centrifuged again at 150,000 *× g* for 10 min at 20 °C. Pellets were resuspended with 1 mL EDTA (ethylene diamine tetra-acetic acid) 100 mM for 10 min at 37°C and centrifuged again at 150,000 *× g* for 10 min at 20 °C to remove EDTA. Pellets were resuspended with 1 mL of acetone for 10 min at RT and centrifuged again at 150,000 *× g* for 10 min at 20 °C. Teichoic acids were eliminated from the pellets by resuspension in 1 mL of 48% hydrofluoric acid and shaking incubation for 48 hours at 4 °C. After four rounds of centrifugation (150,000 *×g* for 10 min at 20 °C), each time washing the pellet with 3 mL cold MilliQ water, the pellets containing PG sacculi were obtained.

### V. cholerae peptidoglycan isolation

For PG isolation from *V. cholerae*, SDS-free pellets from sacculi preparation were resuspended in 100 mM Tris-HCl pH 8, and proteinase K was added to a final concentration of 20 µg/ml and these were incubated for 1 hour at 37 °C. The reaction was stopped by adding 100 µl 10% SDS and boiling for 5 minutes. Sacculi were pelleted by ultracentrifugation for 10 min at 20 °C and 150,000 *×g* three times and each time washed with 3 mL of MiliQ water. Finally, the SDS-free PG sacculi were obtained.

### In vitro activity assays and sample preparation for LC-MS analysis

Protein samples were first purified on a Superdex 200 Increase 10/300 GL column equilibrated with 50 mM Tris-HCl, 300 mM NaCl, 50 mM Tris HCl, pH 7.5. Pellets containing purified PG sacculi were resuspended in hydrolase buffer (100 mM NaPO4, 2 mM NaN3, 100mM NaCl pH 7.5) and Muramidase, Slt70Ec, PrgK_EC_, PrgK_SLT_, PrgK_CHAP_, PrgK_SLT-CHAP_ (including all the variants and combinations) were added to the reaction at a final concentration of 100 µg/ml. When TCEP was used, it was added at a final concentration of 10 mM. All the reactions were incubated at 37 °C overnight (12-16 hours) and subsequently stopped by boiling the reaction for 5 minutes. Once the samples were cooled, 20 μl of borate buffer (0.5 M boric acid adjusted to pH 9 with NaOH) were added (per 100 μL of reaction) to adjust the pH to alkaline. Next, 10 μL of 2 M NaBH4 (a mild reducing agent) was added and the sample was incubated for 30 min at 20 °C. Finally, the pH of the sample was adjusted to 2-3 with 25 % (v/v) orthophosphoric acid and the samples were filtered (0.2 μm pore size) for analysis.

### Peptidoglycan analysis

Ultra Performance Liquid Chromatography (UPLC) was used to perform chromatographic analyses of muropeptides. The UPLC system (Waters) was fitted with an analytical column (BEH C18 column; 130 Å, 1.7 μm, 2.1 mm by 150 mm; Waters, USA) and a trapping cartridge precolumn (SecurityGuard ULTRA Cartridge UHPLC C18 2.1 mm, Phenomenex). Muropeptides were identified using an ACQUITY UPLC UV-visible detector to measure absorbance at 204 nm. Muropeptides were separated over a 15-minute period at a flowrate of 0.25 ml/min using a linear gradient from buffer A (Water + 0.1% (v/v) formic acid) to buffer B (Acetonitrile 100% (v/v) + 0.1% (v/v) formic acid). Muropeptides were quantified from their integrated areas using standards, which were samples at known concentrations. Using a Xevo G2-XS Q-tof system (Waters Corporation, USA), MS-MS/MS analysis was used to confirm the identity of the muropeptides. Positive ionization mode was used to operate the instrument. Muropeptide detection was carried out by mass spectrometry elevated energy (MS^E^) with the following parameters: capillary voltage at 3.0 kV, source temperature at 120 °C, desolvation temperature at 350 °C, sample cone voltage at 40 V, cone gas flow at 100 l/h, desolvation gas flow at 500 l/h, and collision energy (CE): low CE: 6 eV and high CE ramp: 15–40 eV. This allowed for the simultaneous acquisition of precursor and product ion data. A scan speed of 0.25 s/s was used to acquire mass spectra. The scan ranged from 100 to 2000 m/z. Software programs like Masslynx or UNIFI were used for data processing and acquisition (Waters Corp.)

## Results

### X-ray structure and AlphaFold2 modeling

To unravel the function and regulation of PrgK as the cell wall remodeling enzyme, we attempted to overexpress and purify full length PrgK from *E. coli*. Unfortunately, this only gave extremely low yields. Therefore, we directed our efforts towards the extracytoplasmic region (PrgK_EC_), as well as its individual LytM, SLT, and CHAP domains^32^. All these components were successfully produced and purified to homogeneity (Fig. S1). Crystallization trials yielded crystals of the PrgK_LytM_ domain, which were used to solve the LytM structure to 1.5 Å (Fig. 1A, Table 1). In agreement with bioinformatic predictions ^32^, our crystal structure confirms that the active site of LytM is degenerate as it lacks the conserved HXXXD and HXH motifs required for zinc ion coordination at the catalytic site^48,49^(Fig. 1B). This is not the only characterized LytM domain with a degenerate active site. One of the top hits from DALI^50^ and Foldseek^51^ searches with PrgK_LytM_ is the regulatory protein DipM with an r.m.s.d. of 2.68 Å (PDB code: 7QRL) from *C. vibrioides*, which is an activator of the cell wall remodeling amidase AmiC and the lytic transglycosylase SdpA ^52–54^ (Fig. S2A). Sequence alignment of PrgK_LytM_ homologs shows very low sequence identity, with only the region encompassing the HXXXD and HXH motifs being conserved. The His and Asp residues that are important for catalysis are only conserved in active homologs, but not in PrgK_LytM_ and other degenerate homologues (Fig. S2B). In conclusion, the LytM domain of PrgK has a degenerate active site and might play a regulatory role.

**Figure 1.**
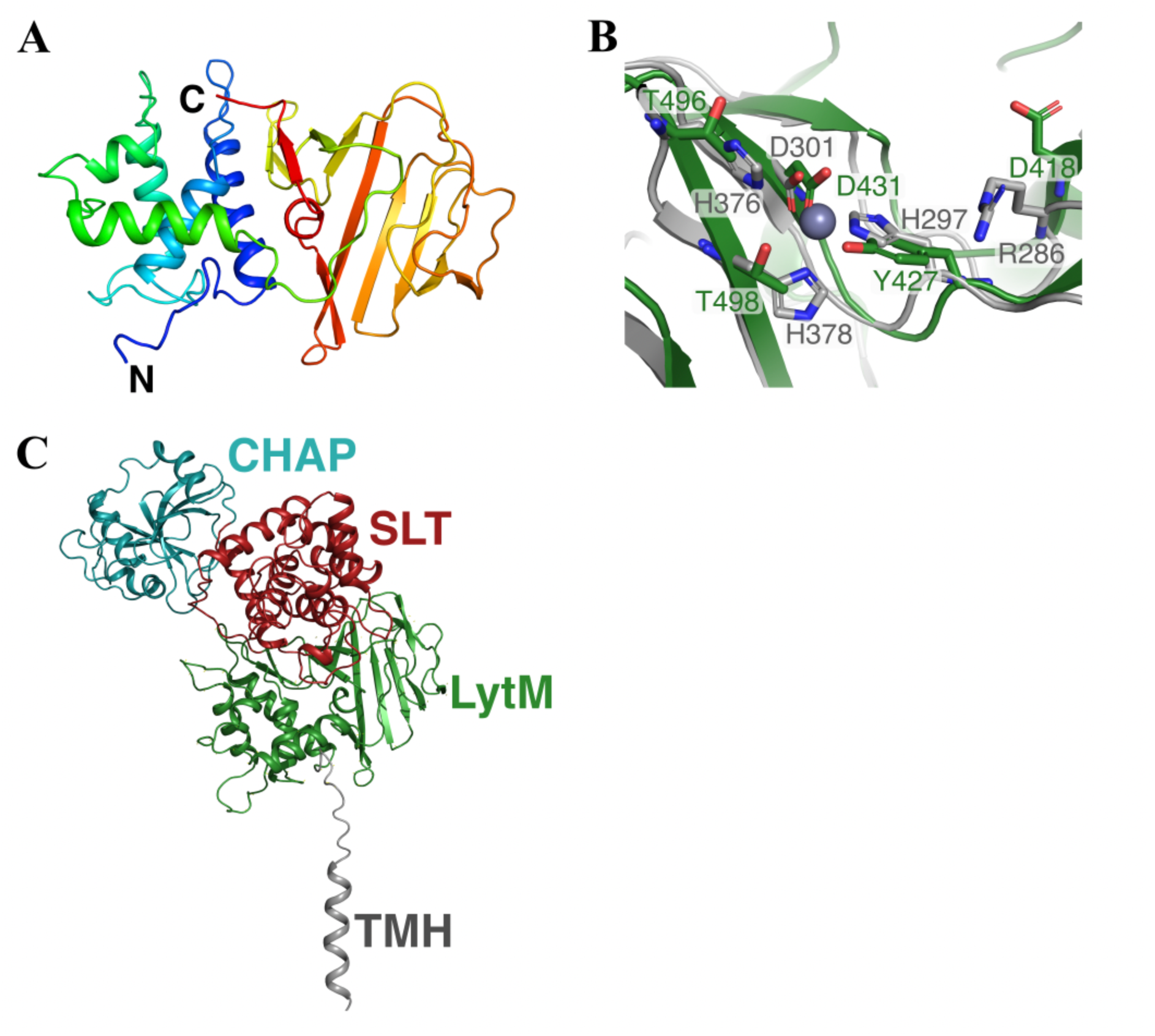
Structure of PrgK. (A) Cartoon representation of the PrgK_LytM_, colored from N-terminal (blue) to C-terminal (red). (B) Comparison of the active site of PrgK_LytM_ (green) and ShyA from *V. cholerae* (gray). Key residues in the active site of ShyA, important for either coordination of the Zn^2+^ (gray sphere) or other activity are shown as sticks, together with the corresponding residues from PrgK_LytM_. (C) AlphaFold2 model of PrgK, showing the transmembrane helix and the three extracellular domains. The intracellular domain is less well predicted and is therefore not shown for clarity.

To obtain a structural model for full-length PrgK, we used AlphaFold2 (AF)^55^. The AF model exhibited low certainty for the intracellular domain but did predict an alpha helix for the transmembrane helix as predicted by TOPCONS ^56^ (residues 254-274) (Fig. 1B). AF confidently predicted the structures of the extracytoplasmic domains, PrgK_EC_, including the relative positioning of the LytM and SLT domains (Fig. 1C & S3A-B). The crystal structure of LytM superimposes well onto the AF model with an r.m.s.d. of 0.43 Å.

Next, we used AF to predict potential interacting partners for PrgK among the other T4SS proteins of pCF10 ^17,19^. This analysis suggested a PrgK:PrgL heterodimer with reasonable confidence. In this AF model, the VirB8-like domain of PrgL (PrgL32-208) ^35^ interacts with the SLT domain of PrgK (Fig. 2A & S4A-B). The analogous T4SS proteins from the conjugative plasmid pIP501, TraG and TraM, have previously been reported to interact^57^, further strengthening this predicted interaction.

**Figure 2.**
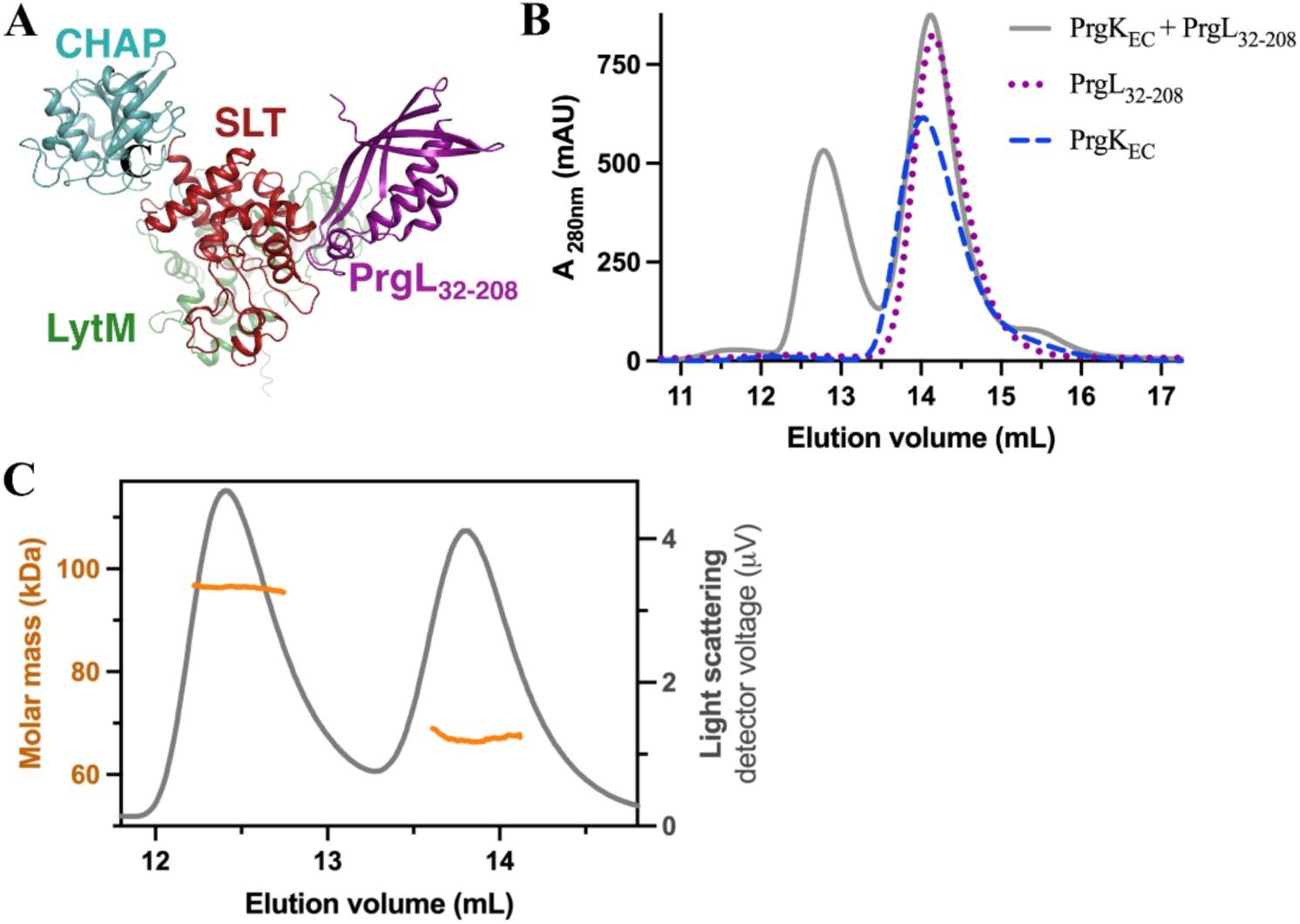
PrgK interacts with PrgL. A) AlphaFold2 model of the PrgK:PrgL heterodimer predicts that PrgL_32-208_ binds to the SLT domain of PrgK. B) Elution profile from size exclusion chromatography of PrgK_EC_, PrgL_32-208_ or a (1:1) mixture of both. The elution profile of the mixture shows an additional peak at an elution volume of 12.7 mL. C) Representative SEC-MALS analysis of the PrgK_EC_ + PrgL_32-208_ (1:1) mixture, with the determined molecular mass indicated as orange lines for both the earlier eluting complex and non-complex (corresponding to the molecular mass of PrgK_EC_ 67.7 kDa) peaks.

### PrgK interacts with PrgL in vitro

To validate the predicted interaction between PrgK and PrgL, we tested whether the purified extracellular domains of the two proteins (PrgK_EC_ and PrgL32-208) could form a complex and co-elute from size exclusion chromatography. PrgL32-208 forms elongated dimers in solution ^35^, which gives it a similar elution volume as PrgK_EC_ (Fig. 2B). However, the elution profiles of PrgK_EC_ and PrgL32-208 mixtures show an additional peak that elutes earlier and that contains both proteins, thus confirming the predicted interaction (Fig. 2A & S4C). To define the stoichiometry of this complex of PrgK_EC_ and PrgL32-208, we performed size-exclusion chromatogram connected to multi-angle light scattering (SEC-MALS). The determined molecular mass of the complex was 91.5 ± 8 kDa (average and standard deviation of 8 independent measurements), which corresponds to a 1:1 complex of the two components (Fig. 2B, PrgK_EC_ is 67.7 kDa and PrgL32-208 is 21.3 kDa).

### Dimerization of PrgK in vitro is dependent on Cys766 in the CHAP domain

During the purifications of PrgK_EC_ we consistently observed a less intense peak that eluted earlier from the SEC. Since the corresponding fraction only contained PrgK_EC_ (Fig. S1A), we speculated that this could correspond to a PrgK_EC_ dimer. This was verified by SEC-MALS experiments, which determined the molecular mass of the two eluting peaks to 64 ± 1 kDa and 144± 4 kDa (average and standard deviation of 3 independent measurements) (Fig. 3C). This corresponds well to the predicted molecular weight of the PrgK_EC_ monomer (67.7 kDa) and dimer (135 kDa).

**Figure 3.**
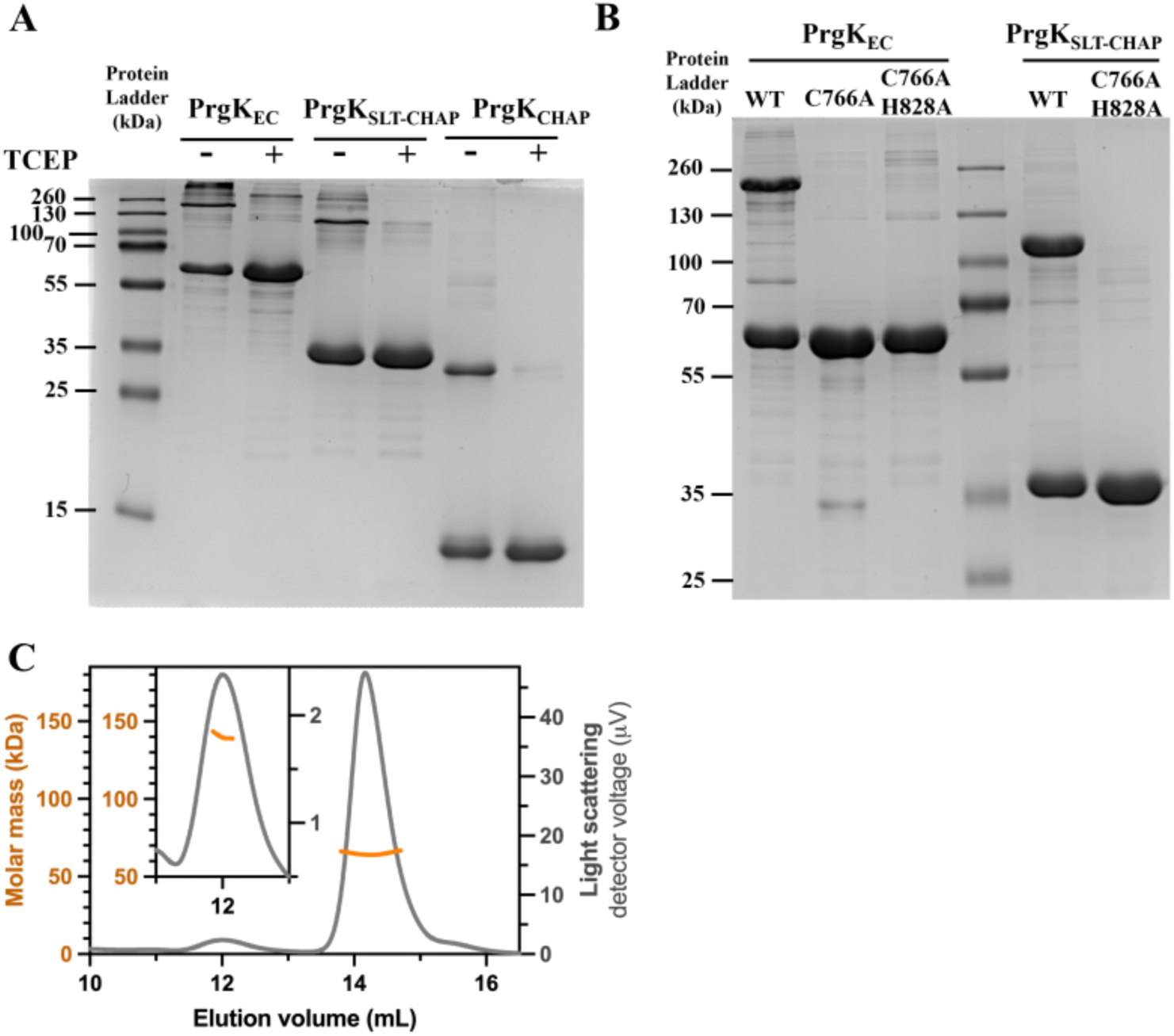
Dimerization of PrgK. A) Non-reducing SDS-PAGE with or without addition of the reducing agent TCEP (to the loaded sample). Bands running at roughly the expected height of a dimer are seen for PrgK_EC_, PrgK_SLT-CHAP_ and PrgK_CHAP_ and in all cases these are significantly decreased upon the addition of TCEP. B) Modifying cysteine 766 in the CHAP domain to an alanine prevents the formation of the dimer band. C) Representative SEC-MALS analysis of PrgK_EC_ (with a theoretical monomeric weight of 67.7 kDa), with the determined molecular mass indicated as orange lines for both the dimeric (shown enlarged in the inset) and monomeric peaks.

To verify whether this partial dimerization was redox dependent, we ran IMAC purified protein on non-reducing SDS-PAGE with or without reducing agents. Without reducing agents, we observe both a monomeric and dimeric band of PrgK_EC_. Importantly, this dimeric band disappears upon the addition of the reducing agent TCEP (Fig. 3A). PrgK_EC_ contains just a single cysteine, located in the predicted active site of the CHAP domain (C766), that could form a disulfide bond. Therefore, PrgKSLT-CHAP and PrgKCHAP were also tested and indeed showed the same redox-dependent oligomerization. To verify that the observed dimerization of PrgK_EC_ is mediated by C766, we created and purified variants where C766 was replaced by an alanine. Indeed, the dimeric band on SDS-PAGE could not be observed for these variants, confirming that C766 can form disulfide bonds and dimerize PrgK (Fig. 3B). Addition of PrgL32-208 did not alter the dimerization properties of PrgK_EC_ in any way (Fig. S4D).

### The SLT domain of PrgK has muramidase activity and it is regulated by the other two EC domains

Periplasmic expression of the extracellular part of PrgK or its individual domains has been described to impair cell growth and lead to cell defects in *E. coli* due to extensive peptidoglycan degradation ^32^. However, there are often mechanisms in place to prevent autolytic enzymes from degrading the cell wall in their native host^58,59^. We therefore wanted to investigate the impact of PrgK expression on *E. faecalis* cells. Full-length PrgK was ectopically overexpressed in the OG1RF strain of *E. faecalis* in the absence of other T4SS proteins. In contrast to the findings in *E. coli*, no major effects were found upon overexpression of PrgK in *E. faecalis*. There were small effects on cell growth and cell viability (Fig. 4A&B), but no phenotypic differences could be observed under scanning electron microscopy (Fig. 4C). We conclude that PrgK overexpression has a small, but significant, impact on *E. faecalis* viability, but does not cause any drastic effect on the morphology of *these* cells.

**Figure 4.**
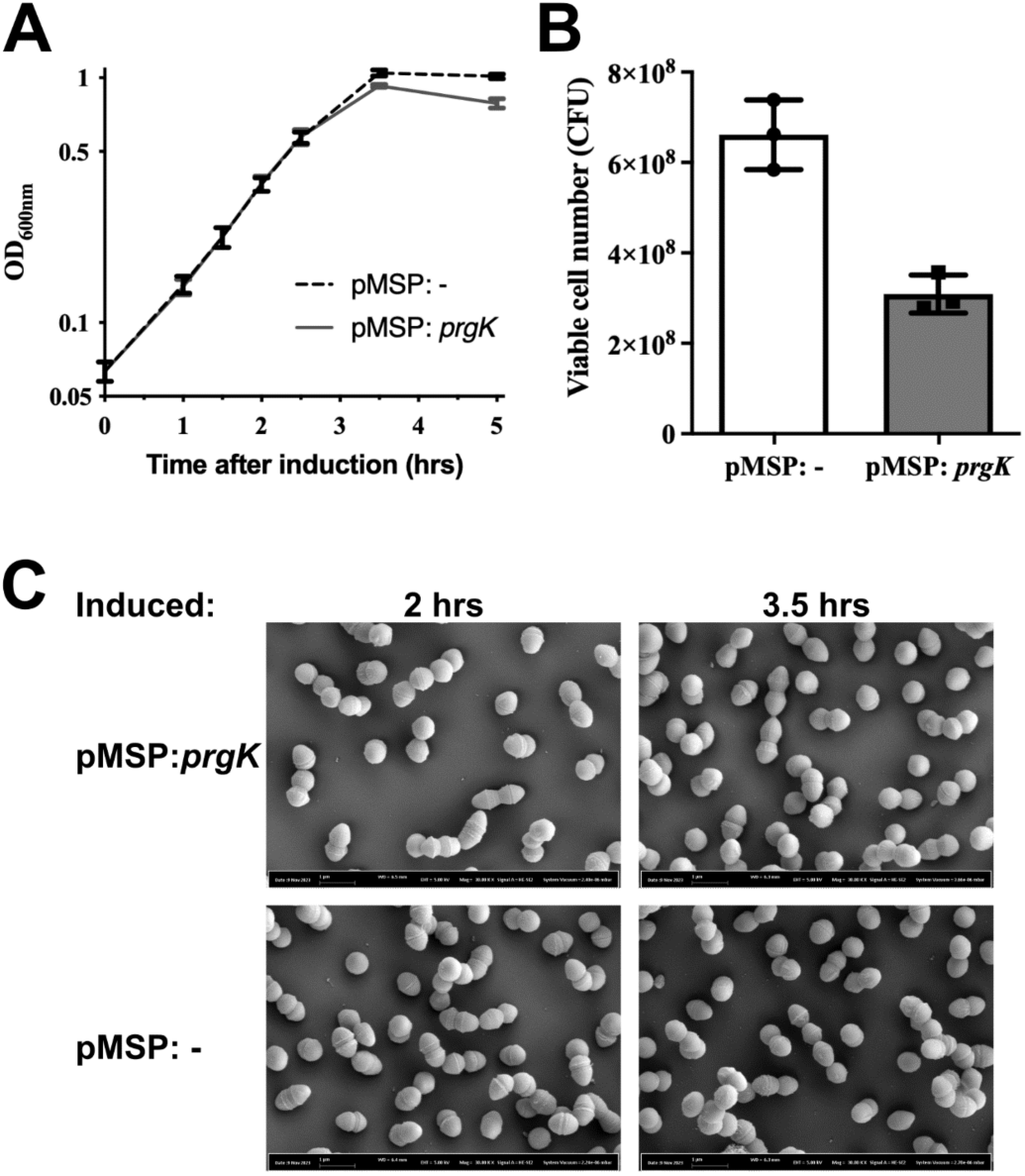
Ectopic expression of PrgK in the absence of other T4SS components. Comparison of *E. faecalis* OG1RF strains harboring a PrgK-expressing plasmid or an empty vector. A) Growth curves of the two strains. Expression was induced at the beginning of incubation (0 hr). B) The viable cell numbers 3.5 hour arer induction. A+B) Data from three independent experiments, error bars indicate standard error of the mean (SEM). C) Representative figures of both strains under scanning electronic microscopy (scanning EM) at 30,000x magnification 2 or 3.5 hrs arer induction.

To characterize the enzymatic and regulatory properties of the PrgK extracellular domains (PrgK_EC_, PrgKSLT, PrgKCHAP, PrgKSLT-CHAP) we assayed their ability to digest the cell wall of *E. faecalis*. Experiments gave similar results when done with sacculi isolated from cCF10-induced *E. faecalis* OG1RF pCF10 as with the OG1RF pCF10:Δ*prgK* strain induced likewise (compare Fig. 5A-B with Fig. S5). This indicates that activity of endogenous PrgK does not significantly alter the cell wall composition after pCF10 is induced by the pheromone cCF10 to initiate T4SS production. Consistent with a previous report ^32^, the SLT domain displayed PG-degrading activity as it digested *E. faecalis* PG into soluble muropeptides (PG fragments). However, in stark contrast with its prediction as a lytic transglycosylase, our UPLC-MS analyses revealed that the PG fragments generated by PrgKSLT were identical to that produced by muramidase (Fig. 5A & S5A). These fragments differ substantially from those generated by a standard SLT protein like the SLT from *E. coli* (Slt70), which is not active against *E. faecalis* PG (Fig. S6A&B).

**Figure 5.**
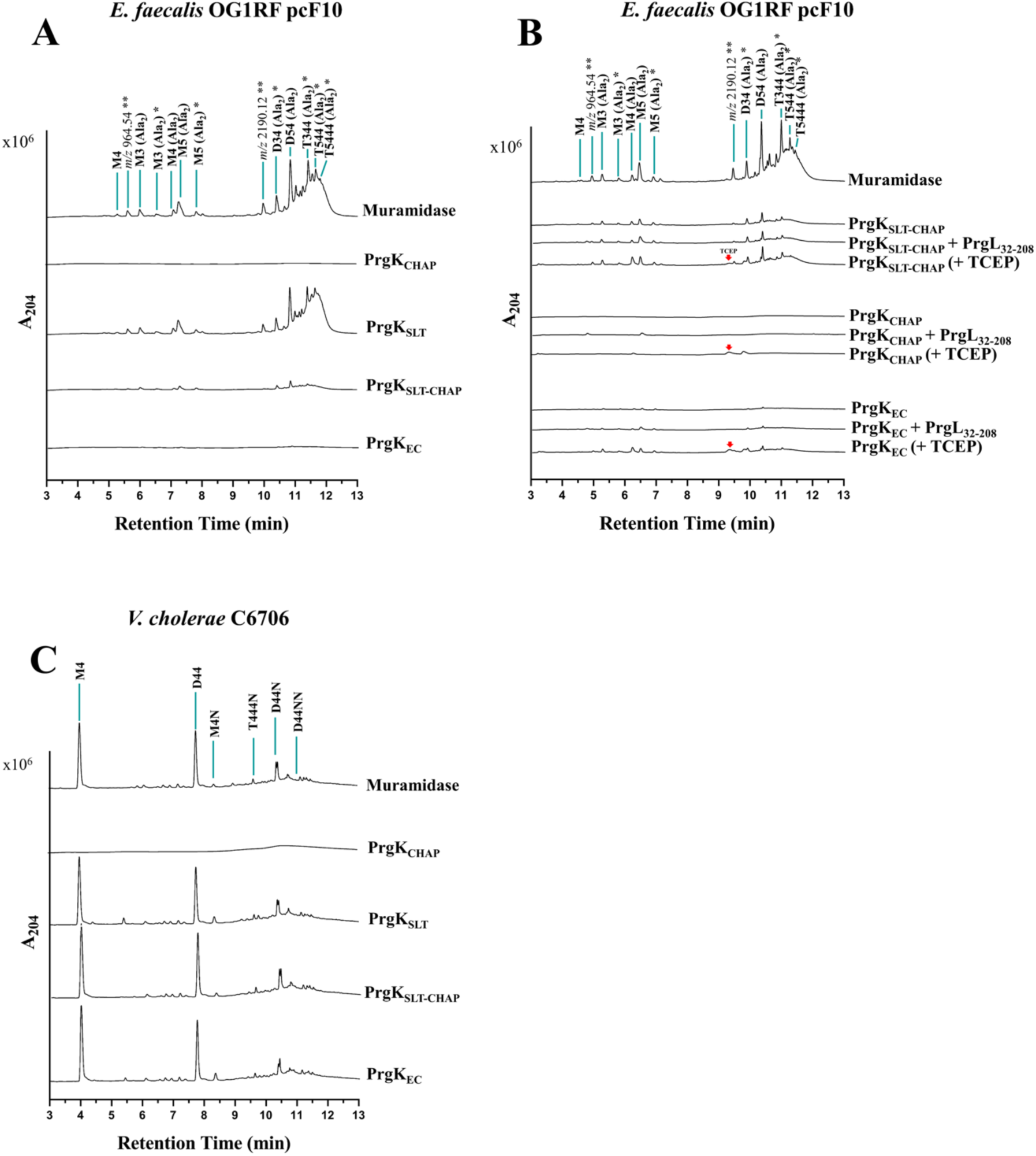
Comparing cell wall muropeptides generated by PrgK domains. A) Chromatograms of muropeptides released following *E. faecalis* OG1RF:pCF10 PG treatment with the indicated PrgK enzymes variants. B) *E. faecalis* OG1RF:pCF10 PG treatment with PrgL_32-208_ or TCEP. C) *V. cholera* C6706 PG treatment with the indicated PrgK enzymes variants. A-C) Identified muropeptides are indicated above their corresponding peak (See table 2 for further information). Unknown muropeptides are marked for their corresponding *m/z* values. Red arrows indicate the peak for TCEP. * Gln/Glu. It is arbitrarily to assign the amide and hydroxyl roles to either pep’de stem. ** Precise structure unknown

Curiously, the activity of PrgKSLT-CHAP was reduced compared to PrgKSLT, and PrgK_EC_ showed an even further reduction (Fig. 5A). To investigate if this reduction in PG degradation was due to the CHAP domain further degrading the products of the SLT domain, we constructed and purified variants of PrgKSLT-CHAP and PrgK_EC_ in which the catalytic Cys766 and His828 residues of the CHAP domain were replaced by Alanines. However, the muropeptide profiles of the PrgK_SLT-CHAP:C766A,H828A_ and PrgK_EC:C766A,H828A_ fractions after incubation with cell-wall extract are similar to their wildtype counterparts (Fig. S6C). In line with this result, the CHAP domain showed no activity against the sacculi (Fig. 5A), as reported earlier ^32^. These experiments therefore exclude the possibility of the hydrolytic activity of CHAP contributing to the diminished muropeptide profiles and instead indicate that CHAP and LytM strongly reduce the activity of the SLT domain in cell wall extracts from *E. faecalis*.

Since we found that PrgK can dimerize in a redox dependent manner, and that it can interact with PrgL, we also performed the activity measurements with PrgL32-208 present or under reducing conditions. According to the data analysis, the hydrolytic activities of PrgKSLT-CHAP and PrgK_EC_ were not altered by PrgL32-208. PrgL32-208 also did not activate PrgKCHAP (Fig. 5B). Addition of the reducing reagent TCEP, to remove any PrgK dimers, showed a small but significant increase in muramidase activity (2-fold increase over no addition, Fig. 5B). This indicates that PrgK dimer formation plays at least a small role in regulating PrgK activity.

Since the effect of PrgK expression in *E. faecalis* cells (Fig. 4) differed substantially from previously reports with PrgK domainsexpression in *E. coli* ^32^, we further wanted to investigate whether the activity of PrgK could be different on cell-wall substrates from G^+^ and G^-^ bacteria. Therefore, we also assayed the ability of PrgKSLT, PrgKCHAP, PrgKSLT-CHAP, and PrgK_EC_ to cleave sacculi from *Vibrio cholerae*, a G^-^ bacteria. This analysis further validated the muramidase-like activity of PrgKSLT. PrgKCHAP did also not show any activity under the tested conditions with this substrate. However, much to our surprise, the activity of PrgKSLT-CHAP and PrgK_EC_ were the same as PrgKSLT, indicating that the downregulation of the activity of the SLT-domain does not occur with Gram-negative cell wall substrates (Fig. 5C).

## Discussion

We have here studied the structure, function, and regulation of PrgK, the cell wall hydrolase of the T4SS from pCF10. Mass spectrometry analysis of cell wall extracts that were enzymatically cleaved by PrgK variants showed unexpectedly that the predicted SLT domain, named after its homology with soluble lytic transglycosylase enzymes, had instead a muramidase activity. Lytic transglycosylases cleave the 1,4-beta-linkages between the MurNAc and GlcNAc sugars with concomitant 1,6-cyclization of the MurNAc, resulting in glycan chains that terminate in a 1,6-anhydromuropeptide at the reducing end ^60–64^. Muramidases cleave the same bond in the PG but do so via a hydrolytic cleavage mechanism using completely different chemistry and thereby producing MurNAc termini without the ring ^65^. We therefore rename the SLT domain in PrgK to MUR (muramidase) domain (Fig. 6A). So far, all other T4SS PG-degrading enzymes have been reported to use proteins with lytic transglycosylase activity ^34^. However, to our knowledge, PrgK is the first T4SS PG-degrading enzyme from the DELTA family that has been characterized in detail.

**Figure 6.**
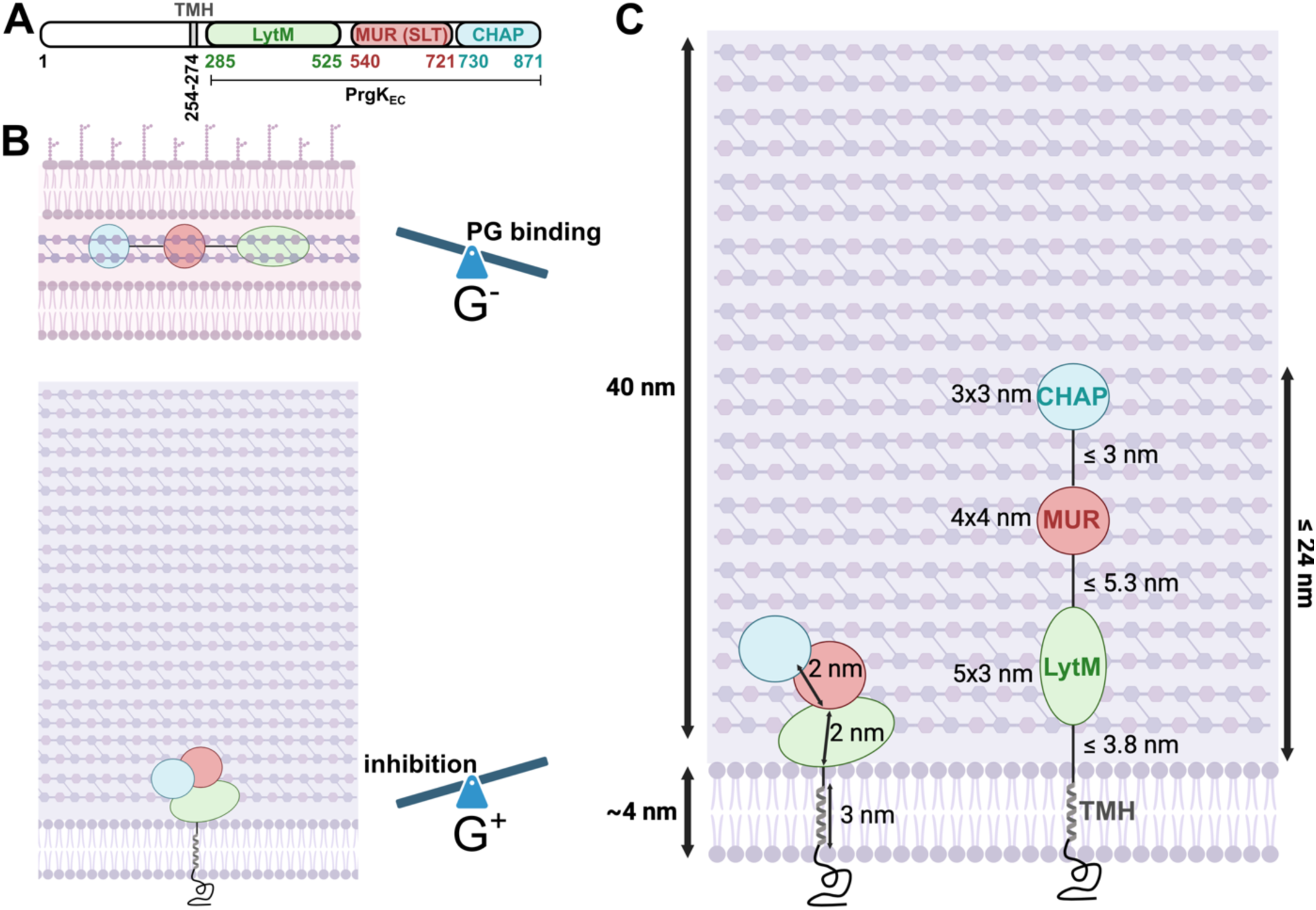
A) Domain organization of PrgK. The formally called SLT domain is renamed MUR domain, because of its murimidase activity. Domain boundaries were predicted by AlphaFold2 and differ slightly from previous predictions. B) Model of PrgK_EC_ PG binding and hydrolysis in G^-^ or G^+^ bacteria. The LytM and CHAP domains can either bind to their PG substrates or interact with the MUR domain (formally termed SLT) thereby blocking its enzyme activity. In G^+^ bacteria, the MUR domain is inhibited by the CHAP and LytM domains as seen in *E. faecalis* cells and cell-wall extracts. In G^-^ bacteria, such as *V. cholerae* or *E. coli*, LytM and CHAP bind stronger to their PG substrate and do not inhibit the MUR domain. C) Maximally extended PrgK (without unfolding the domains), can reach roughly the middle of the *E. faecalis* cell wall.

When we understood that PrgK had muramidase activity, we revisited the AF model of the MUR domain and searched for homologs. Only one structural homolog was found in the PDB by using Foldseek, which is the geneproduct 13 (gp13) cell-wall degrading enzyme of bacteriophage Φ29^66^. gp13 consists of two domains, one N-terminal lysozyme-like domain that is homologous to the MUR domain (RMSD of 1.5 Å), and a C-terminal LytM domain (RMSD to PrgK_LytM_ of 2.4 Å) (Fig. S7). When searching against the AFDB^67^ virtually all top hits are phage tail lysozyme domain containing proteins. It is difficult to determine the evolution of proteins encoded on conjugative plasmids, but it is tempting to speculate that PrgK has picked up the MUR domain (and possibly also the LytM domain) from bacteriophages at some point during its evolution.

Previous work showed that *prgK_lytM_* could not complement a Δ*prgK* strain and the LytM domain was suggested to have a degenerate active site. Our crystal structure confirms this (Fig. 1). The *in vitro* analysis of the CHAP domain also showed virtually no activity, which was surprising given that a PrgK variant lacking the MUR domain, but still having the LytM and CHAP domains could partially rescue a Δ*prgK* strain *in vivo* ^32^. Possibly, PrgK_CHAP_ has a low background level of activity *in vivo* that is sufficient to support conjugation. An analog of PrgK is TraG in the pIP501 T4SS, also from *E. faecalis.* TraG contains an SLT and CHAP domain but lacks a LytM ^33^. The SLT domain of TraG has been indicated to be a lytic transglycosylase, since it is inhibited by specific LT inhibitors ^33^. Both TraGSLT and TraGCHAP domains have also been shown to have relatively limited activities against *E. faecalis* PG in the absence of mutanolysin, and have significantly increased digestion when they react together *in cis* ^33^. These properties of TraG are distinct from PrgK, which has three domains that do not react synergistically. This indicates substantial differences in how the two G^+^ T4SSs modify the cell wall in preparation for conjugation.

Our activity assays showed that PrgK_EC_, containing all three extracellular domains (LytM, MUR and CHAP), only had a very low activity on *E. faecalis* cell-wall extracts (Fig. 5A). The presence of the other two domains thus downregulated the activity of the MUR domain, presumably by sterically blocking the active site. This is analogous to how some other cell-wall degrading enzymes are regulated, like Auto from *Listeria monocytogenes* or ShyA from *V. cholerae* ^58,59^. The predicted AlphaFold2 model of PrgK_EC_ does not indicate that the active site of the MUR domain is blocked by the other domains. However, the PG substrates are large and there is quite some flexibility in the linker regions between the LytM, MUR and CHAP domains that could allow for a reorganization of the three domains and a steric inactivation of the MUR domain. This hypothesis that CHAP and LytM downregulate the activity of the MUR domain is further supported by our <u>phenotype</u> analysis of an *E. faecalis prgK* overexpression strain. Our experiments could only detect a minor effect on cell growth and no changes in the cell-shape upon *prgK* overexpression. If the MUR domain in PrgK had been fully active in these experiments, we would likely have seen a much larger changes as previously reported for *E. coli* expression ^32^.

Interestingly, this downregulation of the MUR domain by the other two domains is not seen when a cell-wall extract from *V. cholerae*, a G^-^ bacteria, is used. There, the MUR domain has the same activity in isolation as in PrgK_EC_. From previous work we know that both the LytM and CHAP domains have the potential to bind peptidoglycan (PG) ^32^. We therefore hypothesize that the LytM and CHAP domains are in an equilibrium between interacting with the MUR domain, thereby sterically blocking its active site, and interacting with the PG (Fig. 6B). In the PG extract from *V. cholerae* or in the periplasm of *E. coli*, this equilibrium could be shifted towards PG interaction, thereby freeing the active site of the MUR and allowing it to bind and digest the PG of these G^-^ bacteria (Fig. 6**B**). The regulation of the MUR domain of PrgK, by the CHAP and LytM domains, that is observed with *E. faecalis* cell-wall extract, is likely important to prevent extra toxicity to the donor cells upon induction of the T4SS. It also seems tuned to the host species, as the regulation is lost on cell wall from G^-^ species. Presumably, PrgK gets activated by another component at an early point after T4SS induction, but it’s not yet clear how this occurs. Analogous to the interaction of TraG and TraM in pIP501 T4SS (Kholer 2017), AlphaFold predicted the interaction of PrgK with PrgL, which we could confirm *in vitro* (Fig. 2). However, PrgL did not alter the activity of PrgK (Fig. 5). We speculate that PrgL binds PrgK to localize it to the T4SS channel during channel assembly. This is in line with the previous hypothesis that PrgL, as other VirB8-like proteins, acts as a scaffold during the biogenesis of the T4SS ^35^.

During the purification of PrgK_EC_, we noted that a small amount of the protein eluted earlier from SEC. This fraction was confirmed to be a redox dependent dimer of PrgK_EC_, that was formed by disulfide bonds between the single cysteine, C766, from two CHAP domains. C766 is an integral part of the active site of the CHAP domain and dimer formation via this residue will therefore likely shut down any potential activity that the CHAP domain has. Our cell wall activity data could not detect any activity for the CHAP domain. This absence of enzymatic activity makes sense if the function of the CHAP domain is not hydrolysis, but instead regulation and dimer formation.

A large question in the field of G^+^ T4SS is how the substrate gets transported into the recipient cell. No pili exist in these systems, but instead cell wall hydrolases have been suggested to form a discrete lesion in the cell walls of both cells, basically providing a physical tunnel of sorts between the mating pair. However, structural analysis of PrgK reveals that even if the protein would be completely extended (without unfolding the individual domains), it would only reach roughly the middle of the cell wall of the *E. faecalis* donor cell (Fig. 6C). Thus, we deem it very unlikely that PrgK could create a defined substrate tunnel between the donor and recipient bacteria. Instead, we propose that the main function of PrgK is to create enough space in the cell wall of the donor cell to allow for channel assembly. This means that it remains unclear how the substrate transfers with a high efficiency from the donor cell, over two cell wall barriers and finally entering the recipient cell by somehow passing over its membrane. Answering this question will be of great interest in future research.

To conclude, we here show that PrgK, an essential cell wall degrading enzyme of the pCF10 T4SS, has muramidase activity instead of the lytic transglycosylase activity that has been shown for other T4SS cell wall degrading enzymes. Only the MUR domain has measurable activity *in vitro*, while the other two extracellular domains of PrgK instead have a regulatory function. This results in the autoregulation of PrgK hydrolysis activity, which likely is important to prevent toxicity for the donor cell. Finally, we have also shown that PrgK interacts with PrgL, which sheds further light into the assembly mechanisms of Gram-positive T4SSs.

## Acknowledgments

We acknowledge MAX IV Laboratory for time on Beamline BioMax under Proposal 20180236. Research conducted at MAX IV, a Swedish national user facility, is supported by the Swedish Research council under contract 2018-07152, the Swedish Governmental Agency for Innovation Systems under contract 2018-04969, and Formas under contract 2019-02496. We acknowledge the European Synchrotron Radiation Facility (ESRF) for provision of synchrotron radiation facilities under proposal number mx2326 and we would like to thank A. McCarthy for assistance and support in using beamline ID30B. This work was supported by grants from the Swedish Research Council (2016-03599), Knut and Alice Wallenberg Foundation, Kempestiftelserna (SMK-1762 & SMK-1869) to R.P-A.B and from the Swedish Research Council (2018-02823 and 2018-05882), Umeå University, the Knut and Alice Wallenberg Foundation and the Kempe Foundation (SMK2062) to F.C. G.T. is supported by an EC postdoctoral fellowship supported by the Knut and Alice Wallenberg Foundation.

## CRediT Author Contributions

Wei-Sheng Sun: Conceptualization, Investigation, Writing – Original Draft, Writing – Revision. Gabriel Torrens: Investigation, Writing – Revision. Kai Ehrenbolger: Investigation, Writing – Original Draft. Josy ter Beek: Conceptualization, Investigation, Writing – Original Draft, Writing – Revision. Felipe Cava: Supervision, Funding acquisition, Writing – Revision. Ronnie P-A Berntsson: Conceptualization, Writing – Original Draft, Writing – Revision, Supervision, Funding acquisition.

---

**Supplementary figure 1.**
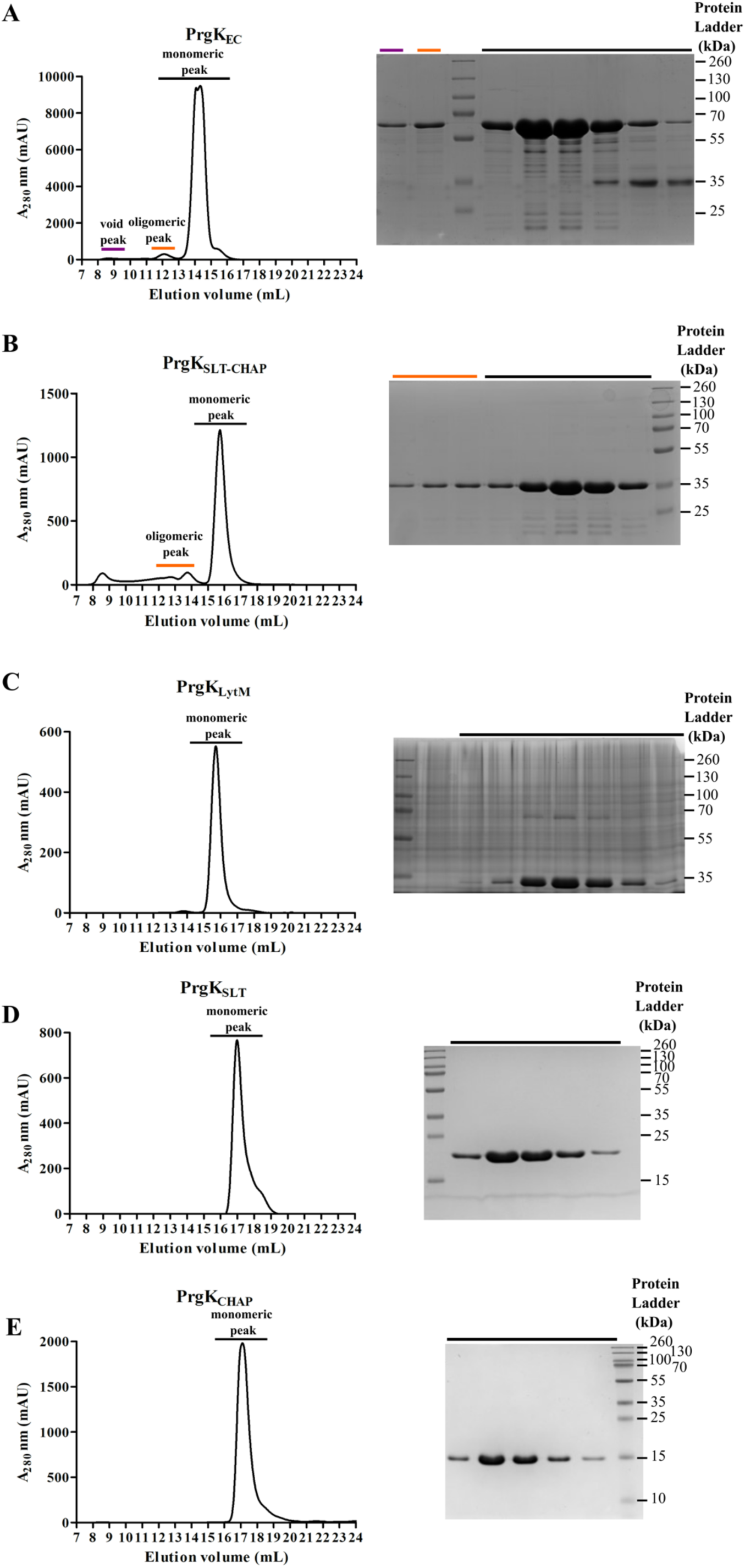
Representative elution profiles from size exclusion chromatography (ler panels) and corresponding Coomassie-blue stained SDS-PAGE (right panels) of PrgK variants. Additional peaks other than the main peaks are marked with the same color in the ler and right panels: A) PrgK_EC_, consisting of residue 273-871, B) PrgK_SLT-CHAP_, consisting of residue 530-871, C) PrgK_LytM_, consisting of residue 273-529, D) PrgK_SLT_, consisting of residue 539-723, E) PrgK_CHAP_, consisting of residue 723-871.

**Supplementary figure 2.**
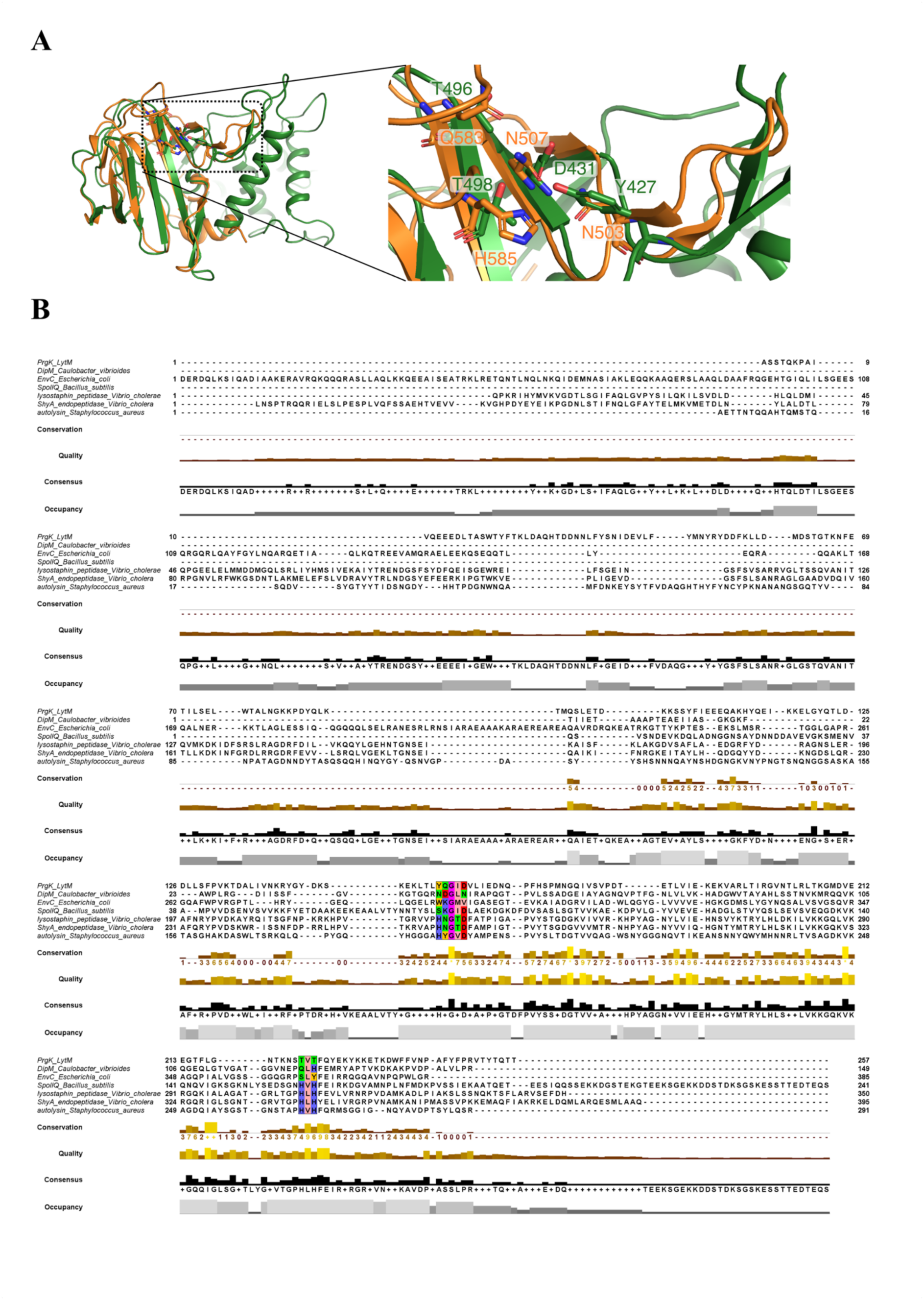
Comparison of PrgK_LytM_ to homologs. A) Ler panel: Structure superimposition of *C. vibrioides* DipM (orange) to PrgK_LytM_ (green). The degenerate catalytic centers of DipM and PrgK_LytM_ are highlighted in the dashed box and enlarged in the right panel, in which the residues corresponding to the HX_3_D and HXH motifs are displayed as sticks. B) Primary sequence alignment of PrgK_LytM_ homologues. The HX_3_D and HXH motifs that are essential for catalysis are highlighted in colored squares. Homologous proteins include catalytically degenerate ones: *C. vibrioides* DipM (7qrl, 461-609), *E. coli* EnvC (6tpi, 35-419), and *B. subtilis* SpIIQ (3tuf, 43-283); catalytically active ones are exemplified by lysostaphin peptidase from *Vibrio cholerae* (2gu1, 63-412), *V. cholerae* ShyA endopeptidase (6u2a, 36-430), and *S. aureus* autolysin LytM (1qwy, 26-316). Consensus sequence is displayed as logos below the individual sequences. Number on the ler and right are scales, not the actual residue numbers.

**Supplementary figure 3.**
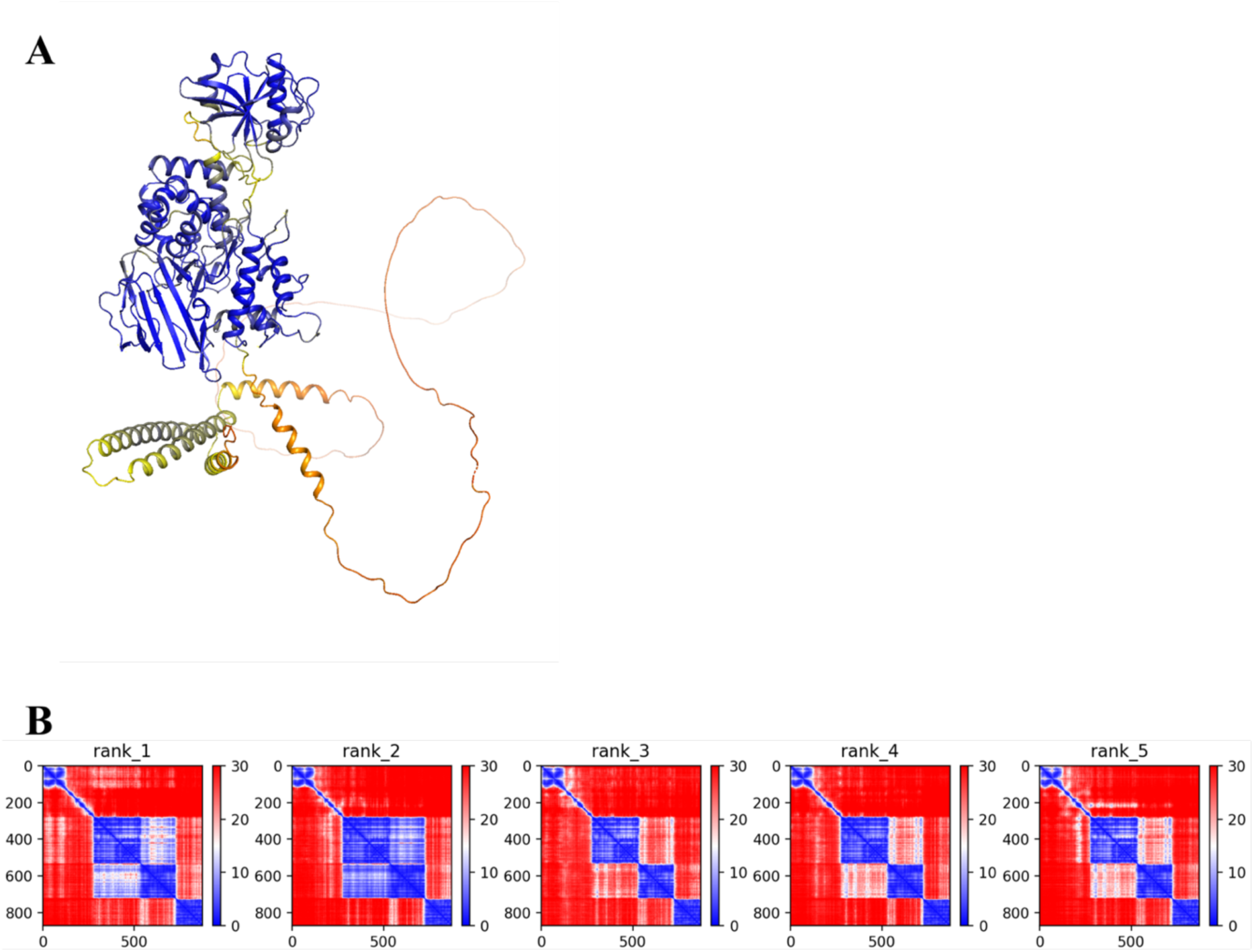
AlphaFold2 model of full length PrgK. A) pLDDT colored full length PrgK, including the intracellular domain. Colors range from red (low quality) to blue (high quality). B) PAE plots of the top five models generated by AlphaFold2. The model is reasonably confident in the placement of the LytM and SLT domain in relation to each other, but not the remaining domains.

**Supplementary figure 4.**
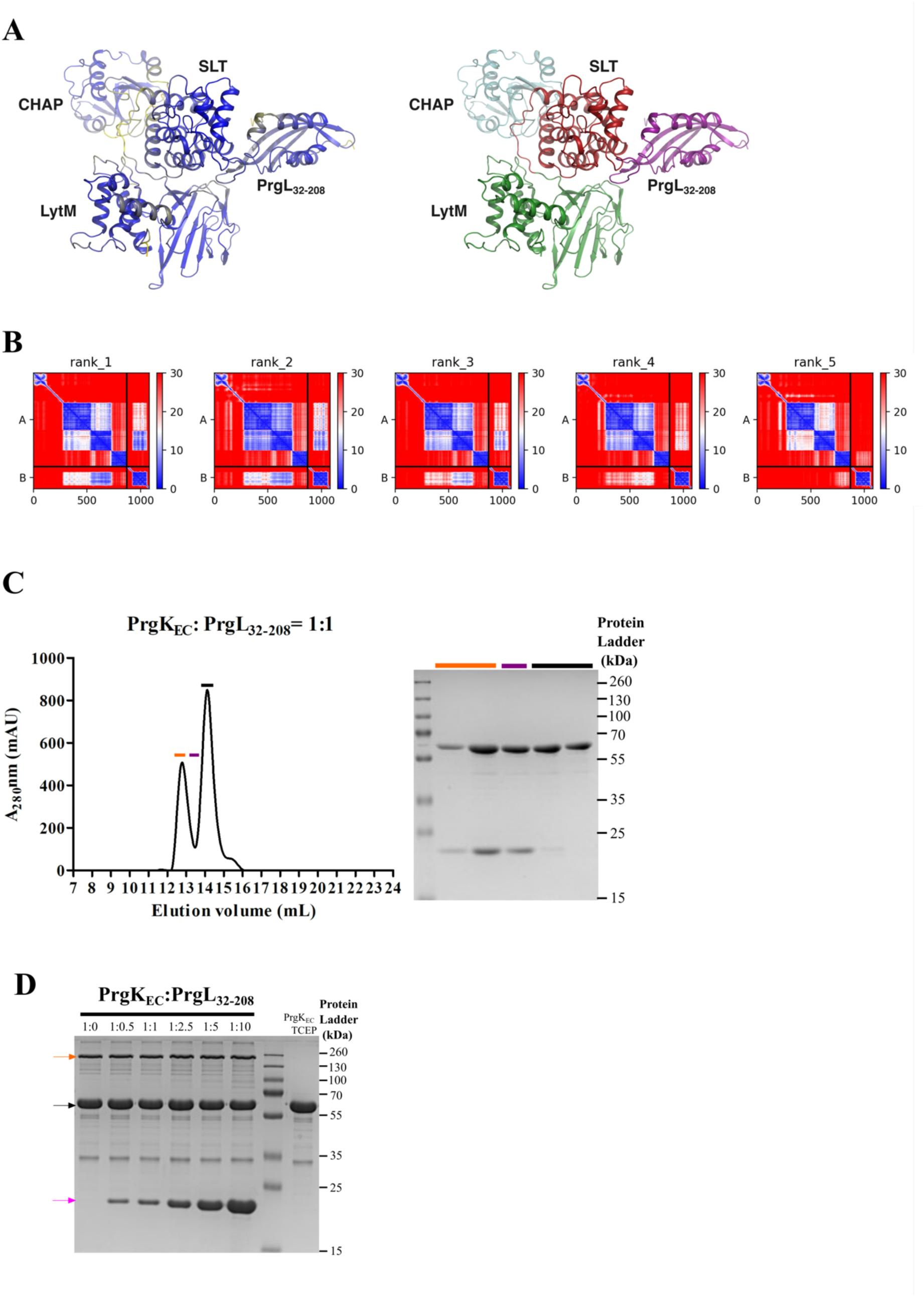
PrgK-PrgL protein-protein interaction. A) AlphaFold2 multimer prediction of protein complex formed between PrgK and PrgL. Ler: colored by pLDDT, right: colored by domains. B) PAE plots of the top five AlphaFold2 models of PrgK:PrgL. C) Ler: representative size exclusion chromatography elution profile (Superdex 200 Increase 10/300 GL column) of PrgK_EC_ and PrgL_32-208_ mixture. Right: SDS-PAGE of fractions covering the first peak (orange), second peak (black), and the interval (purple). D) *In vitro* dimerization of PrgK_EC_ is not altered in the presence of PrgL_32-208_. The protein bands of dimeric PrgK_EC_ (indicated by orange arrow) and monomeric PrgK_EC_ (indicated by black arrow) in the presence of increasing amounts of PrgL_32-208_ (indicated by purple arrows). The molar ratios of PrgK_EC_:PrgL_32-208_ are indicated above each lane. PrgK_EC_ incubated with TCEP, which reduces the dimer to monomers, is shown as a control (final lane).

**Supplementary figure 5.**
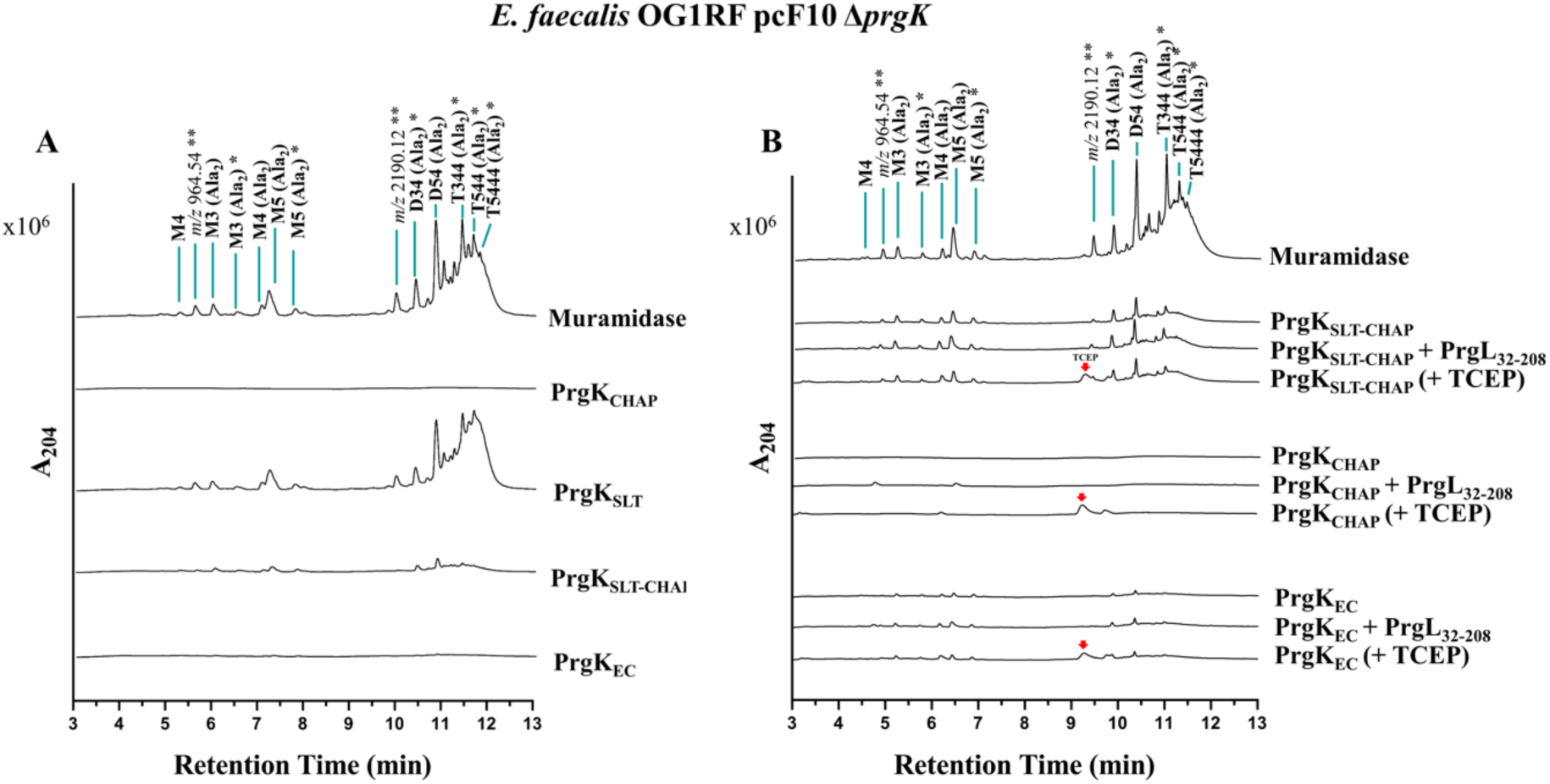
Comparing cell wall muropeptides generated by PrgK domains. Chromatograms of muropeptides released following *E. faecalis* OG1RF:pCF10Δ*prgK* PG treatment with the indicated PrgK enzymes variants (A) and with addition of PrgL_32-208_ or TCEP (B). Identified muropeptides are indicated above their corresponding peak (See table 2 for further information). Unknown muropeptides are marked for their corresponding *m/z* values. Red arrows indicate the peak for TCEP. * Gln/Glu. It is arbitrarily to assign the amide and hydroxyl roles to either pep’de stem. ** Precise structure unknown

**Supplementary figure 6.**
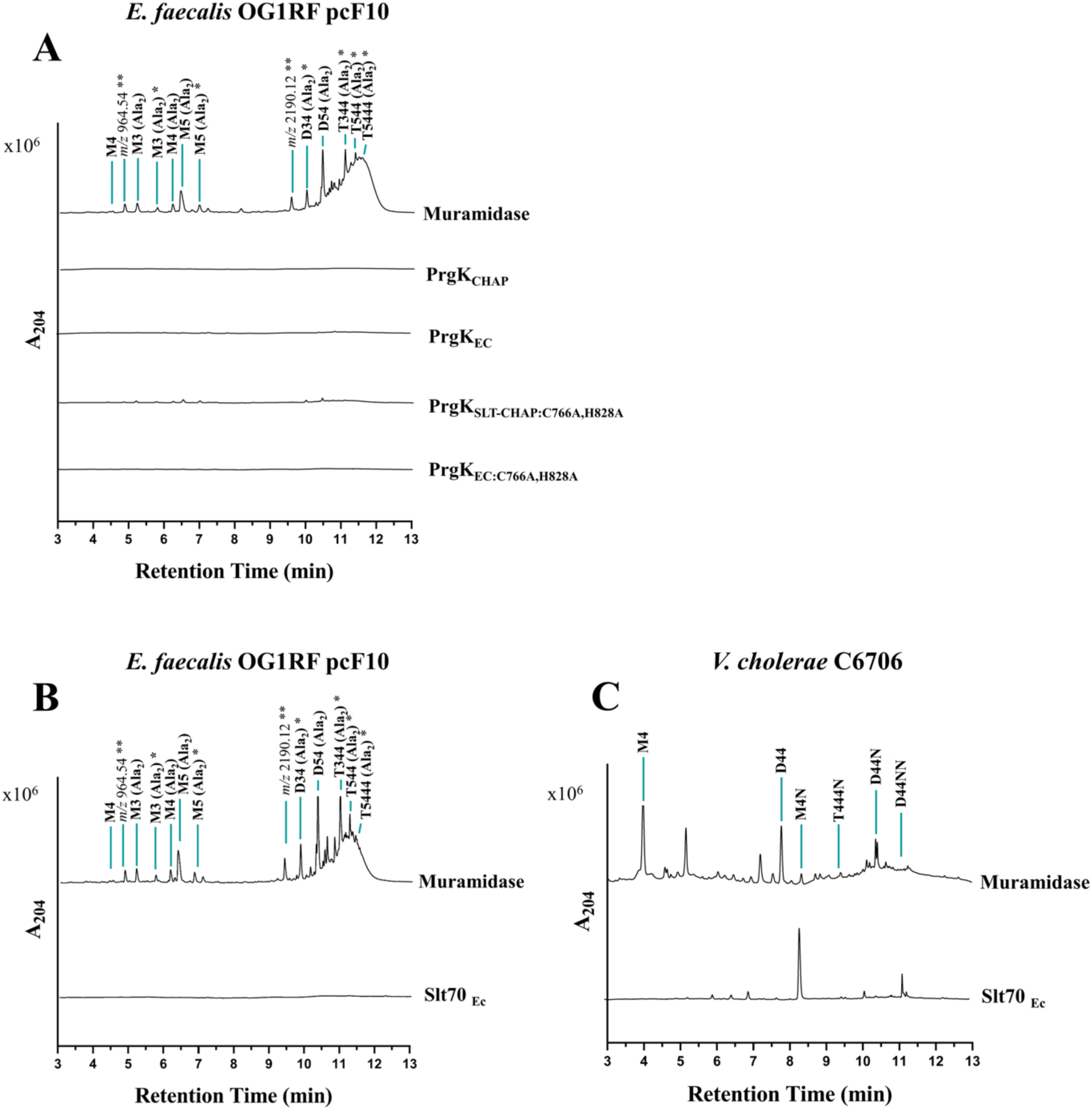
Activity of PrgK_SLT_. A) Mutating the active site residues C766 and H828 to alanines in the CHAP domain does not change the activity of PrgK B) Slt70 from *E. coli* does not function on sacculi from *E. faecalis*. C) Slt70 from *E. coli* does cleave sacculi isolated from *V. cholerae*. * Gln/Glu. It is arbitrarily to assign the amide and hydroxyl roles to either peptide stem. ** Precise structure unknown

**Supplementary Figure 7.**
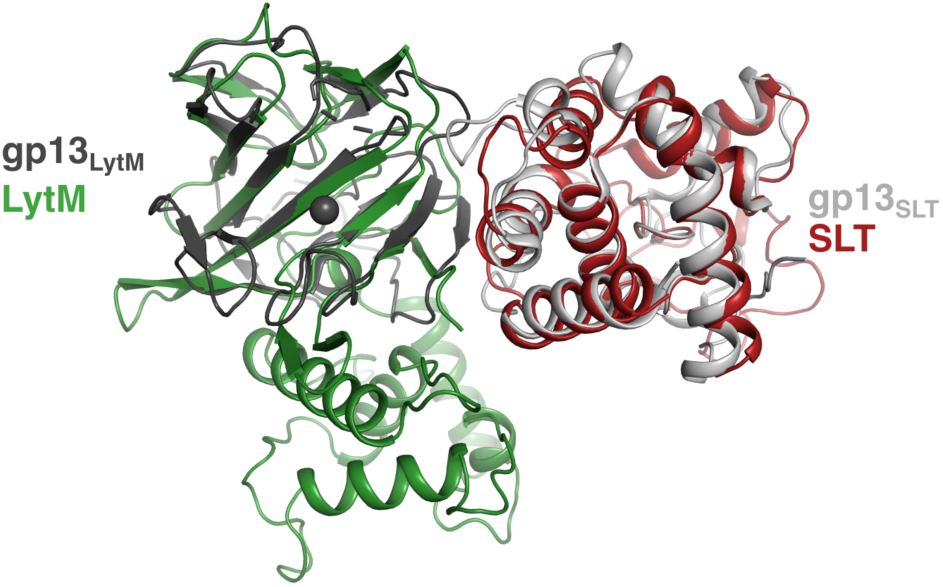
Comparison of PrgK_LytM+SLT_ AlphaFold2 model with the crystallized cell-wall degrading enzyme gp13 of bacteriophage Φ29 (PDB code:3CSQ). The SLT domain of gp13 (light gray) and the LytM domain of gp13 (dark gray) superimpose well onto PrgK_SLT_ and PrgK_LytM_ (RMSD of 1.5 Å and 2.4 Å), respectively (the gp13 domains were superimposed individually onto PrgK, and do not represent the correct orientation in the gp13 crystal structure). In contrast to PrgK_LytM_, the gp13_LytM_ domain has a functional active site, and has a Zn^2+^ bound. (shown as sphere)

**Table S1.**
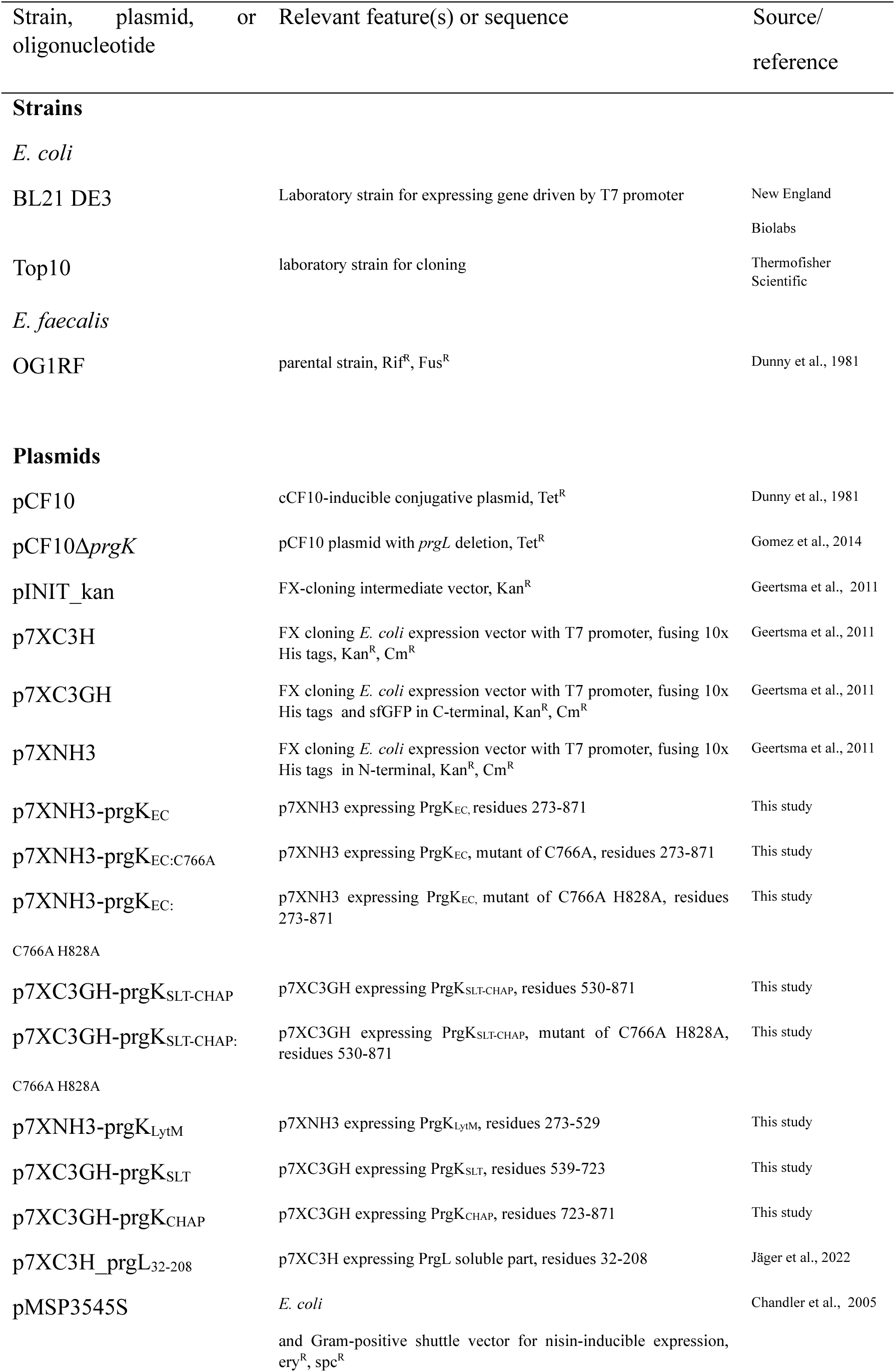
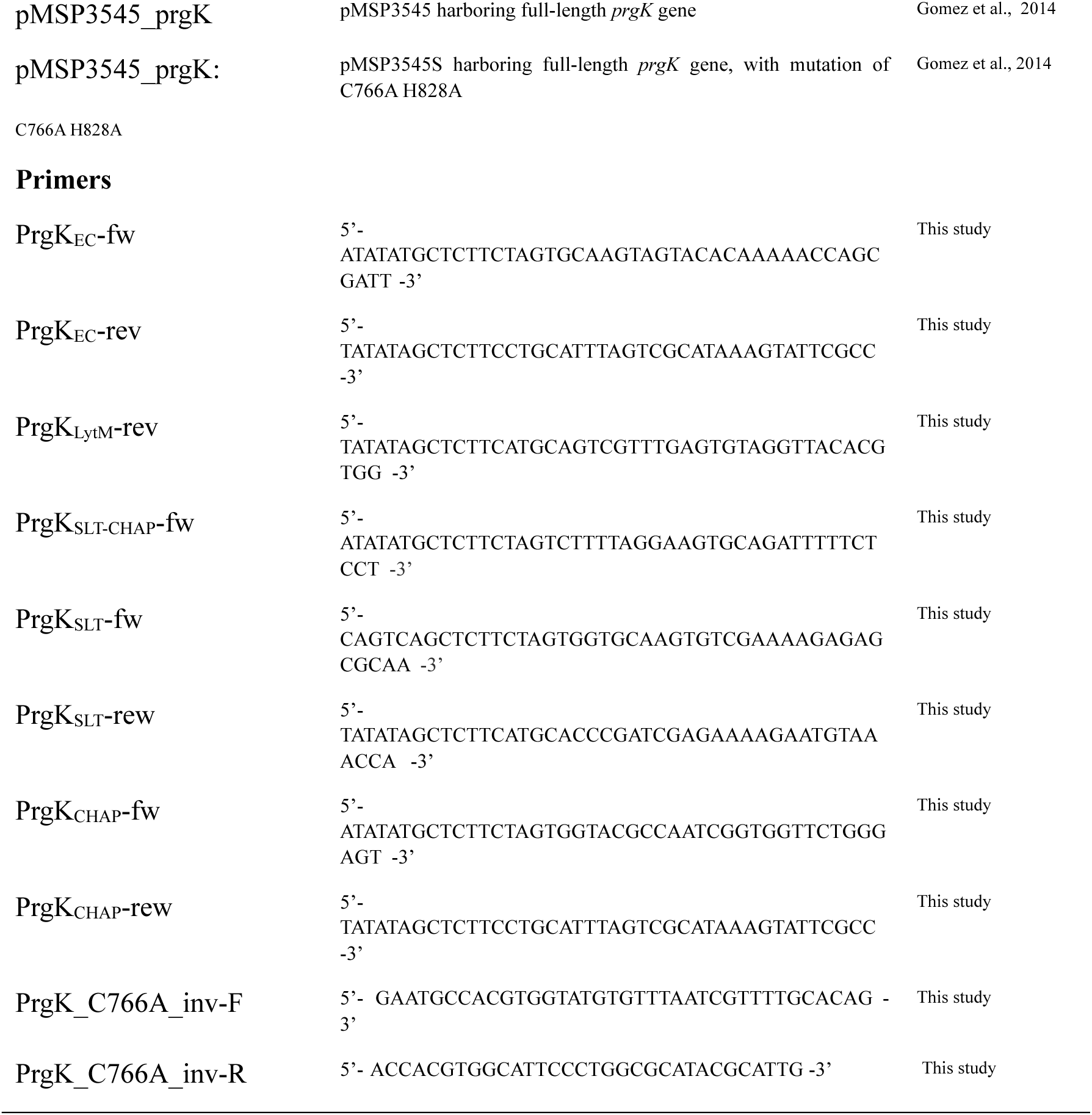
Table of bacterial strains, plasmids and nucleotides used.

**Table S2.** Molecular mass and proposed structure of muropeptides from *E. faecalis*.

| Identity | Proposed Structure | Observed *m/z* | Observations |
| --- | --- | --- | --- |
| M4 | GlcNAc-MurNAc-L-Ala-D-iGln-L-Lys-D-Ala | 897.44 | [M+H]<sup>+</sup> |
| ND | Precise structure unknown | 964.54 | [M+H]<sup>+</sup> |
| M3 (Ala<sub>2</sub>) | GlcNAc-MurNAc-L-Ala-D-iGln-L-Lys(-L-Ala-L-Ala) | 968.48 | [M+H]<sup>+</sup> |
| M3 (Ala<sub>2</sub>)<sup>*</sup> | GlcNAc-MurNAc-L-Ala-D-iGlu-L-Lys(-L-Ala-L-Ala) | 969.46 | [M+H]<sup>+</sup> |
| M4 (Ala<sub>2</sub>) | GlcNAc-MurNAc-L-Ala-D-iGln-L-Lys(-L-Ala-L-Ala)-D-Ala | 1039.52 | [M+H]<sup>+</sup> |
| M5 (Ala<sub>2</sub>) | GlcNAc-MurNAc-L-Ala-D-iGln-L-Lys(-L-Ala-L-Ala)-D-Ala-D-Ala | 1110.55 | [M+H]<sup>+</sup> |
| M5 (Ala<sub>2</sub>)<sup>*</sup> | GlcNAc-MurNAc-L-Ala-D-iGlu-L-Lys(-L-Ala-L-Ala)-D-Ala-D-Ala | 1111.53 | [M+H]<sup>+</sup> |
| ND | Precise structure unknown | 2190.12 | [M+H]<sup>+</sup> |
| D34 (Ala<sub>2</sub>)<sup>*</sup> | M3 (Ala<sub>2</sub>) - M4 (Ala<sub>2</sub>) | 996.99 | [M+2H]<sup>2+</sup> |
| D54 (Ala<sub>2</sub>) | M5 (Ala<sub>2</sub>) - M4 (Ala<sub>2</sub>) | 1067.03 | [M+2H]<sup>2+</sup> |
| T344 (Ala<sub>2</sub>)<sup>*</sup> | M3 (Ala<sub>2</sub>) - M4 (Ala<sub>2</sub>) - M4 (Ala<sub>2</sub>) | 1005.17 | [M+3H]<sup>3+</sup> |
| T544 (Ala<sub>2</sub>)<sup>*</sup> | M5 (Ala<sub>2</sub>) - M4 (Ala<sub>2</sub>) - M4 (Ala<sub>2</sub>) | 1052.85 | [M+3H]<sup>3+</sup> |
| T5444 (Ala<sub>2</sub>)<sup>*</sup> | M5 (Ala<sub>2</sub>) - M4 (Ala<sub>2</sub>) - M4 (Ala<sub>2</sub>) - M4 (Ala<sub>2</sub>) | 1045.27 | [M+4H]<sup>4+</sup> |

GlcNAc-MurNAc: N-acetylglucosamine and N-acetylmuramic acid; Ala, Alanine; Gln, Glutamine; Glu, Glutamic acid; Lys, Lysine.
\* It is arbitrary to assign the amide and hydroxyl functions to either peptide stem.

## Notes

### Competing Interest Statement

The authors have declared no competing interest.

## References

1. Garsin, D. A., et al. Pathogenesis and Models of Enterococcal Infection. Enterococci: From Commensals to Leading Causes of Drug Resistant Infection (2014).

2. Lebreton, F. et al. Tracing the Enterococci from Paleozoic Origins to the Hospital. Cell 169, (2017).

3. Sparo, M., Delpech, G. & Allende, N. G. Impact on public health of the spread of high-level resistance to gentamicin and vancomycin in enterococci. Frontiers in Microbiology vol. 9 Preprint at 10.3389/fmicb.2018.03073 (2018).

4. Ch’ng, J. H., Chong, K. K. L., Lam, L. N., Wong, J. J. & Kline, K. A. Biofilm-associated infection by enterococci. Nature Reviews Microbiology vol. 17 Preprint at 10.1038/s41579-018-0107-z (2019).

5. Dunny, G. M., Hancock, L. E. & Shankar, N. Enterococcal Biofilm Structure and Role in Colonization and Disease. Enterococci: From Commensals to Leading Causes of Drug Resistant Infection (2014).

6. Kristich, C. J., Rice, L. B. & Arias, C. A. Enterococcal Infection—Treatment and Antibiotic Resistance. Enterococci: From Commensals to Leading Causes of Drug Resistant Infection (2014).

7. European Centre for Disease Prevention and Control. Antimicrobial resistance surveillance in Europe 2016. Annual Report of the European Antimicrobial Resistance Surveillance Network (EARS-Net). European Centre for Disease Prevention and Control (2017).

8. Hu, B., Khara, P. & Christie, P. J. Structural bases for F plasmid conjugation and F pilus biogenesis in Escherichia coli. Proc Natl Acad Sci U S A 116, 14222–14227 (2019).

9. Khara, P., Song, L., Christie, P. J. & Hu, B. In Situ Visualization of the pKM101-Encoded Type IV Secretion System Reveals a Highly Symmetric ATPase Energy Center. mBio 12, (2021).

10. Hu, B. et al. In Situ Molecular Architecture of the Helicobacter pylori Cag Type IV Secretion System. mBio 10, e00849–19 (2019).

11. Macé, K. et al. Cryo-EM structure of a type IV secretion system. Nature 607, 191–196 (2022).

12. Chung, J. M. et al. Structure of the Helicobacter pylori Cag Type IV secretion system. Elife 8, (2019).

13. Costa, T. R. D., Patkowski, J. B., Macé, K., Christie, P. J. & Waksman, G. Structural and functional diversity of type IV secretion systems. Nature Reviews Microbiology Preprint at 10.1038/s41579-023-00974-3 (2023).

14. Kohler, V., Keller, W. & Grohmann, E. Regulation of Gram-Positive Conjugation. Front Microbiol 10, 1134 (2019).

15. Grohmann, E., Christie, P. J., Waksman, G. & Backert, S. Type IV secretion in Gram-negative and Gram-positive bacteria. Mol Microbiol 107, 455–471 (2018).

16. Wisniewski, J. A. & Rood, J. I. The Tcp conjugation system of Clostridium perfringens. Plasmid 91, 28–36 (2017).

17. Lassinang, L. et al. Enterococcal PrgU Provides Additional Regulation of Pheromone-Inducible Conjugative Plasmids. mSphere 6, e00264–21 (2021).

18. Dunny, G. M. & Berntsson, R. P.-A. Enterococcal sex pheromones: Evolutionary pathways to complex, two-signal systems. J Bacteriol 198, 1556–1562 (2016).

19. Dunny, G. M. Enterococcal Sex Pheromones: Signaling, Social Behavior, and Evolution. Annu Rev Genet 47, 457–482 (2013).

20. Sun, W.-S. et al. Structural foundation for the role of enterococcal PrgB in conjugation, biofilm formation and virulence. Elife 12, RP84427 (2023).

21. SchmiU, A. et al. PrgB promotes aggregation, biofilm formation, and conjugation through DNA binding and compaction. Mol Microbiol 109, 291–305 (2018).

22. SchmiU, A. et al. Enterococcal PrgA Extends Far Outside the Cell and Provides Surface Exclusion to Protect against Unwanted Conjugation. J Mol Biol 432, 5681–5695 (2020).

23. Rehman, S. et al. Enterococcal PcfF is a ribbon-helix-helix protein that recruits the relaxase PcfG through binding and bending of the oriT sequence. Front Microbiol 10, (2019).

24. Chen, Y. et al. Enterococcus faecalis PcfC, a spatially localized substrate receptor for type IV secretion of the pCF10 transfer intermediate. J Appl Microbiol 190, 3632–3645 (2008).

25. Li, F. et al. Enterococcus faecalis PrgJ, a VirB4-like ATPase, Mediates pCF10 Conjugative Transfer Through Substrate Binding. J Appl Microbiol 194, 4041–4051 (2012).

26. Huang, K. C., Mukhopadhyay, R., Wen, B., Gitai, Z. & Wingreen, N. S. Cell shape and cell-wall organization in Gram-negative bacteria. Proc Natl Acad Sci U S A 105, (2008).

27. Silhavy, T. J., Kahne, D. & Walker, S. The Bacterial Cell Envelope1 T. J. Silhavy, D. Kahne and S. Walker, . Cold Spring Harb Perspect Biol 2, (2010).

28. Chang, J. D., Wallace, A. G., Foster, E. E. & Kim, S. J. Peptidoglycan Compositional Analysis of Enterococcus faecalis Biofilm by Stable Isotope Labeling by Amino Acids in a Bacterial Culture. Biochemistry 57, (2018).

29. Llosa, M., Zupan, J., Baron, C. & Zambryski, P. The N- and C-terminal portions of the Agrobacterium VirB1 protein independently enhance tumorigenesis. J Bacteriol 182, (2000).

30. Zupan, J., Hackworth, C. A., Aguilar, J., Ward, D. & Zambryski, P. VirB1* promotes T-pilus formation in the vir-type IV secretion system of Agrobacterium tumefaciens. J Bacteriol 189, (2007).

31. Sgro, G. G. et al. Bacteria-killing type IV secretion systems. Frontiers in Microbiology vol. 10 Preprint at 10.3389/fmicb.2019.01078 (2019).

32. Laverde Gomez, J. A., BhaUy, M. & Christie, P. J. PrgK, a Multidomain Peptidoglycan Hydrolase, Is Essential for Conjugative Transfer of the Pheromone-Responsive Plasmid pCF10. J Appl Microbiol 196, 527–539 (2014).

33. Arends, K. et al. TraG Encoded by the pIP501 Type IV Secretion System Is a Two-Domain Peptidoglycan-Degrading Enzyme Essential for Conjugative Transfer. J Appl Microbiol 195, 4436–4444 (2013).

34. Goessweiner-Mohr, N., Arends, K., Keller, W. & Grohmann, E. Conjugation in Gram-Positive Bacteria. Microbiol Spectr 2, (2014).

35. Jäger, F., Lamy, A., Sun, W.-S., Guerini, N. & Berntsson, R. P.-A. Structure of the enterococcal T4SS protein PrgL reveals unique dimerization interface in the VirB8 protein family. Structure 30, 876–885.e5 (2022).

36. Bae, T., Kozlowicz, B. & Dunny, G. M. Two targets in pCF10 DNA for PrgX binding: their role in production of Qa and prgX mRNA and in regulation of pheromone-inducible conjugation. J Mol Biol 315, 995–1007 (2002).

37. Bryan, E. M., Bae, T., Kleerebezem, M. & Dunny, G. M. Improved vectors for nisin-controlled expression in gram-positive bacteria. Plasmid 44, 183–190 (2000).

38. Källberg, M. et al. Template-based protein structure modeling using the RaptorX web server. Nat Protoc 7, (2012).

39. Kelley, L. A., Mezulis, S., Yates, C. M., Wass, M. N. & Sternberg, M. J. E. The Phyre2 web portal for protein modeling, prediction and analysis. Nat Protoc 10, 845–858 (2015).

40. Geertsma, E. R. & Dutzler, R. A versatile and eficient high-throughput cloning tool for structural biology. Biochemistry 50, 3272–3278 (2011).

41. Kabsch, W. XDS. Acta Crystallographica Section D 66, 125–132 (2010).

42. McCoy, A. J. et al. Phaser crystallographic sorware. J Appl Crystallogr 40, 658–674 (2007).

43. Terwilliger, T. C. et al. Iterative model building, structure refinement and density modification with the PHENIX AutoBuild wizard. 64, 61–69 (2008).

44. Emsley, P., Lohkamp, B., ScoU, W. G. & Cowtan, K. Features and development of Coot. 66, 486–501 (2010).

45. Chen, V. B. et al. MolProbity: all-atom structure validation for macromolecular crystallography. 66, 12–21 (2010).

46. Mirdita, M. et al. ColabFold: making protein folding accessible to all. Nat Methods 19, (2022).

47. Alvarez, L., Hernandez, S. B., De Pedro, M. A. & Cava, F. Ultra-sensitive, high-resolution liquid chromatography methods for the high-throughput quantitative analysis of bacterial cell wall chemistry and structure. in Methods in Molecular Biology vol. 1440 (2016).

48. Firczuk, M., Mucha, A. & Bochtler, M. Crystal Structures of Active LytM. J Mol Biol 354, 578– 590 (2005).

49. Ragumani, S., Kumaran, D., Burley, S. K. & Swaminathan, S. Crystal structure of a putative lysostaphin peptidase from Vibrio cholerae. Proteins: Structure, Function and Genetics 72, 1096–1103 (2008).

50. Holm, L. & Laakso, L. M. Dali server update. Nucleic Acids Res 44, W351–W355 (2016).

51. van Kempen, M. et al. Fast and accurate protein structure search with Foldseek. Nat Biotechnol (2023) doi:10.1038/s41587-023-01773-0.

52. Izquierdo-Martinez, A. et al. DipM controls multiple autolysins and mediates a regulatory feedback loop promoting cell constriction in Caulobacter crescentus. Nat Commun 14, (2023).

53. Peters, N. T. et al. Structure-function analysis of the LytM domain of EnvC, an activator of cell wall remodelling at the Escherichia coli division site. Mol Microbiol 89, 690–701 (2013).

54. Yang, D. C., Tan, K., Joachimiak, A. & Bernhardt, T. G. A conformational switch controls cell wall-remodelling enzymes required for bacterial cell division. Mol Microbiol 85, 768–781 (2012).

55. Jumper, J. et al. Highly accurate protein structure prediction with AlphaFold. Nature 596, (2021).

56. Tsirigos, K. D., Peters, C., Shu, N., Käll, L. & Elofsson, A. The TOPCONS web server for consensus prediction of membrane protein topology and signal peptides. Nucleic Acids Res 43, (2015).

57. Kohler, V. et al. Conjugative type IV secretion in Gram-positive pathogens: TraG, a lytic transglycosylase and endopeptidase, interacts with translocation channel protein TraM. Plasmid 91, 9–18 (2017).

58. Shin, J. H. et al. Structural basis of peptidoglycan endopeptidase regulation. Proc Natl Acad Sci U S A 117, (2020).

59. Bublitz, M. et al. Structural basis for autoinhibition and activation of Auto, a virulence-associated peptidoglycan hydrolase of Listeria monocytogenes. Mol Microbiol 71, (2009).

60. Byun, B. et al. Mechanism of the Escherichia coli MltE lytic transglycosylase, the cell-wall-penetrating enzyme for Type VI secretion system assembly. Sci Rep 8, (2018).

61. Dik, D. A., Marous, D. R., Fisher, J. F. & Mobashery, S. Lytic transglycosylases: concinnity in concision of the bacterial cell wall. Critical Reviews in Biochemistry and Molecular Biology vol. 52 Preprint at 10.1080/10409238.2017.1337705 (2017).

62. Hoeltje, J. V., Mirelman, D., Sharon, N. & Schwarz, U. Novel type of murein transglycosylase in Escherichia coli. J Bacteriol 124, (1975).

63. Williams, A. H. et al. A step-by-step in crystallo guide to bond cleavage and 1,6-anhydro-sugar product synthesis by a peptidoglycan-degrading lytic transglycosylase. Journal of Biological Chemistry 293, (2018).

64. Thunnissen, A. W. H., Isaacs, N. W. & Dijkstral, B. W. The catalytic domain of a bacterial lytic transglycosylase defines a novel class of lysozymes. Proteins: Structure, Function, and Bioinformatics 22, (1995).

65. Herlihey, F. A. & Clarke, A. J. Controlling autolysis during flagella insertion in gram-negative bacteria. in Advances in Experimental Medicine and Biology vol. 925 (2017).

66. Xiang, Y. et al. Crystal and cryoEM structural studies of a cell wall degrading enzyme in the bacteriophage φ29 tail. Proc Natl Acad Sci U S A 105, (2008).

67. Varadi, M. et al. AlphaFold Protein Structure Database: Massively expanding the structural coverage of protein-sequence space with high-accuracy models. Nucleic Acids Res 50, (2022).

